# Dynamic nucleosome organization after fertilization reveals regulatory factors for mouse zygotic genome activation

**DOI:** 10.1101/2022.03.03.482916

**Authors:** Chenfei Wang, Chuan Chen, Xiaoyu Liu, Chong Li, Qiu Wu, Xiaolan Chen, Lingyue Yang, Xiaochen Kou, Yanhong Zhao, Hong Wang, Yawei Gao, Yong Zhang, Shaorong Gao

## Abstract

Chromatin remodeling is essential for epigenome reprogramming after fertilization. However, the underlying mechanisms of chromatin remodeling remain to be explored. Here, we investigated the dynamic changes in nucleosome occupancy and positioning in pronucleus-stage zygotes using ultra low-input MNase-seq. We observed distinct features of inheritance and reconstruction of nucleosome position in both paternal and maternal genomes. Genome-wide *de novo* nucleosome occupancy in the paternal genome was observed as early as 1 hour after the injection of sperm into ooplasm. The nucleosome positioning pattern was continually rebuilt to form nucleosome depletion regions (NDRs) at promoters and transcription factor (TF) binding sites with differential dynamics in paternal and maternal genomes. NDRs formed more quickly on the promoters of genes involved in zygotic genome activation (ZGA), and this formation is closely connected with histone acetylation, but not transcription elongation or DNA replication. Importantly, we found that NDR establishment on the binding motifs of specific TFs might be associated with their potential pioneer functions in ZGA. Further investigations suggested that the predicted factors MLX and RFX1 played important roles in regulating minor and major ZGA, respectively. Our data not only elucidate the nucleosome positioning dynamics in both male and female pronuclei following fertilization, but also provide an efficient method for identifying key transcription regulators during development.

## Introduction

After the sperm penetrates into the oocyte, the chromatin of highly specialized male and female pronuclei undergoes remarkable reprogramming, which supports the transition from meiosis to mitosis and the reactivation of transcription in embryos^1^. In mammals, the oocyte finishes its second meiotic division and exits the meiosis to form the female pronucleus, and the sperm undergoes chromatin de-condensation and protamine-histone replacement to form the male pronucleus^1, 2^. Recently, the dynamics of maternal-to-zygotic transition after fertilization in mammals have been characterized according to specific epigenetic features and the transcription machinery, including DNA modification, histone modification, high-order chromatin architecture and RNA polymerase II (Pol II) binding^3–8^. These studies provide new insights into the chromatin remodeling after fertilization and valuable resources for investigating the mechanisms of transcriptional activation in early embryos. However, the detailed process of protamine-to-histone transition in the paternal genome as well as the dynamics of chromatin state in the maternal genome shortly after fertilization remains to be elucidated.

Nucleosomes are the basic units of chromatin structure and serve multiple cellular functions; they have a compact structure which inhibits the access of sequence-specific proteins. The genome-wide pattern of nucleosome positioning is determined by the combination of DNA sequence, ATP-dependent chromatin remodeling enzymes, and transcription factors (TFs)^9, 10^. In the eukaryotic genome, nucleosome-depleted regions (NDRs) are observed at regulatory elements, including many promoters, enhancers, and terminator regions^10–12^. RNA Pol II passaging results in upstream trafficking of histone proteins and the formation of a typical NDR at the transcription start site (TSS)^13^, and the nucleosome unit downstream or upstream of the NDR is referred to as the +1 nucleosome or −1 nucleosome, respectively. Meanwhile, nucleosomes also serve as barriers for RNA Pol II elongation and impact the gene activation logic and expression noise^14^. In recent years, different regulatory models of zygotic genome activation (ZGA) after fertilization were proposed, in which tight temporal coupling between chromatin reorganization and minor/major waves of ZGA was widely discussed but still under debate^7^. The recently published landscapes of RNA Pol II binding in mouse embryos reveal that Pol II is preferentially loaded to CG-rich promoters and accessible distal regions in one-cell embryo, and the loading of Poll II to future gene targets occurs earlier before genome activation^8^. However, the detailed patterns of chromatin remodeling especially on the view of nucleosome positioning short after fertilization remain unclear. Moreover, whether Poll II or certain pioneer transcription factors coordinate with other chromatin remodelers to participate in NDR formation and how this process affects downstream gene expression as well as ZGA at the early stages remain a long-standing question.

Here, we optimized micrococcal nuclease digestion-based high-throughput sequencing (MNase-seq) to elucidate the nucleosome organization dynamics during the first 12 hours after fertilization. We investigated the dynamics of nucleosome establishment and re-positioning in the male and female pronuclei, respectively. Importantly, through integrative analyses of the NDR formation pattern on TF motifs, we identified novel regulators of zygotic genome activation for mouse early embryos.

## Results

### Mapping nucleosome remodeling in mouse pronuclei

To study the chromatin remodeling of parental pronuclei (PN), we developed an ultra-low-input MNase-seq (ULI-MNase-seq) method using a single tube for micrococcal nuclease digestion^15^ and subsequent library construction (Fig. S1a). We first validated the nucleosome profiles of mouse embryonic stem cells (mESCs) using 100 cells, 5 cells, or a single cell per reaction. Reassuringly, lengths of the mapped reads were enriched at approximately 147 bp (Fig. S1b), corresponding with the mono-nucleosome size^16^. In addition, the genome-wide profiles from 5 or 100 mESCs were highly consistent with the published data from bulk samples^17^ (Fig. S1c-e). Furthermore, we observed precisely positioned +1 and −1 nucleosomes as well as clear NDRs at TSSs, enhancers, and CTCF-binding sites in 5-cell and 100-cell samples (Fig. S1f-h). All these results demonstrated that our ULI-MNase-seq procedures could detect the genome-wide position of nucleosomes with as few as 5 cells. In addition, we developed a computational workflow called NEPTUNE (i**N**tegrat**E**d **P**ipeline **T**o analyze **U**ltra-low-input **N**ucleosome s**E**quencing data), which could analyze ULI-MNase-seq data systematically (Fig. S2).

To avoid heterogeneity due to differences in the timing of fertilization, we used intracytoplasmic sperm injection (ICSI) to approximately establish the same starting time point for 5-8 embryos per group (Fig. 1a). We observed the formation and dynamic changes in parental pronuclei after ICSI (Fig. S3a) and injected H2B-RFP mRNA into oocytes before ICSI to detect the protamine-histone replacement shortly after fertilization. We found that the H2B-RFP signal appeared as early as 1 hour post-fertilization (hpf) (Fig. 1b), and the parental pronuclei were formed at approximately 3 hpf (Fig. S3a), which was corresponding to the PN1 stage^18^. The pronuclei further developed to the PN3 stage at 6 hpf and reached each other at 12 hpf. Based on these observations, we collected parental pronuclei using micromanipulation at different time points (from 0.5 to 12 hpf) and performed ULI-MNase-seq to detect the chromatin state of the pronuclei from formation to fusion (Fig. 1a). Meanwhile, we performed round spermatid injection (ROSI) as a negative control for histone-to-protamine transition in the paternal genome, as the round spermatid (RS) possesses nucleosome-based chromatin instead of protamines^19^. In these experiments, 10-15 pronuclei were used for each reaction. We applied NEPTUNE on these ULI-MNase-seq datasets, and as shown, the biological replicates presented high reproducibility, with exceptions of sperm and 0.5-hpf male PN samples (Fig. S3b-c). The length distribution of nucleosome reads showed a preference for approximately 147 bp in the oocyte, RS, and most PN samples (Fig. S3d). However, a large fraction of short DNA fragments (5-50 bp) was observed in sperm and 0.5-hpf male PN, but they were not observed in the oocyte, RS, or other PN samples (Fig. S3e), which was consistent with the spermatid-specific DNA packaging structures detected by MNase digestion in previous studies^15, 20^. To exclude the possibility of overdigestion^15^, we applied MNase digestion with different concentrations and durations to the sperm samples and confirmed that short fragments were observed in all conditions (Fig. S3f). In agreement with the previous discovery revealed by DNA FISH^20^, the distribution preference of short (5-50 bp) and long (120-180 bp) fragments were distinct from each other (Fig. S3g). The proportion of short fragments remained high in 0.5-hpf male PN, which was dramatically decreased in 1-hpf male PN, suggesting that the sperm-specific DNA packaging structures were largely remodeled in the paternal genome; the remodeling occurred in conjunction with or after the removal of protamines, which was detected at 25-35 min post-fertilization by imaging^21^.

**Fig. 1.**
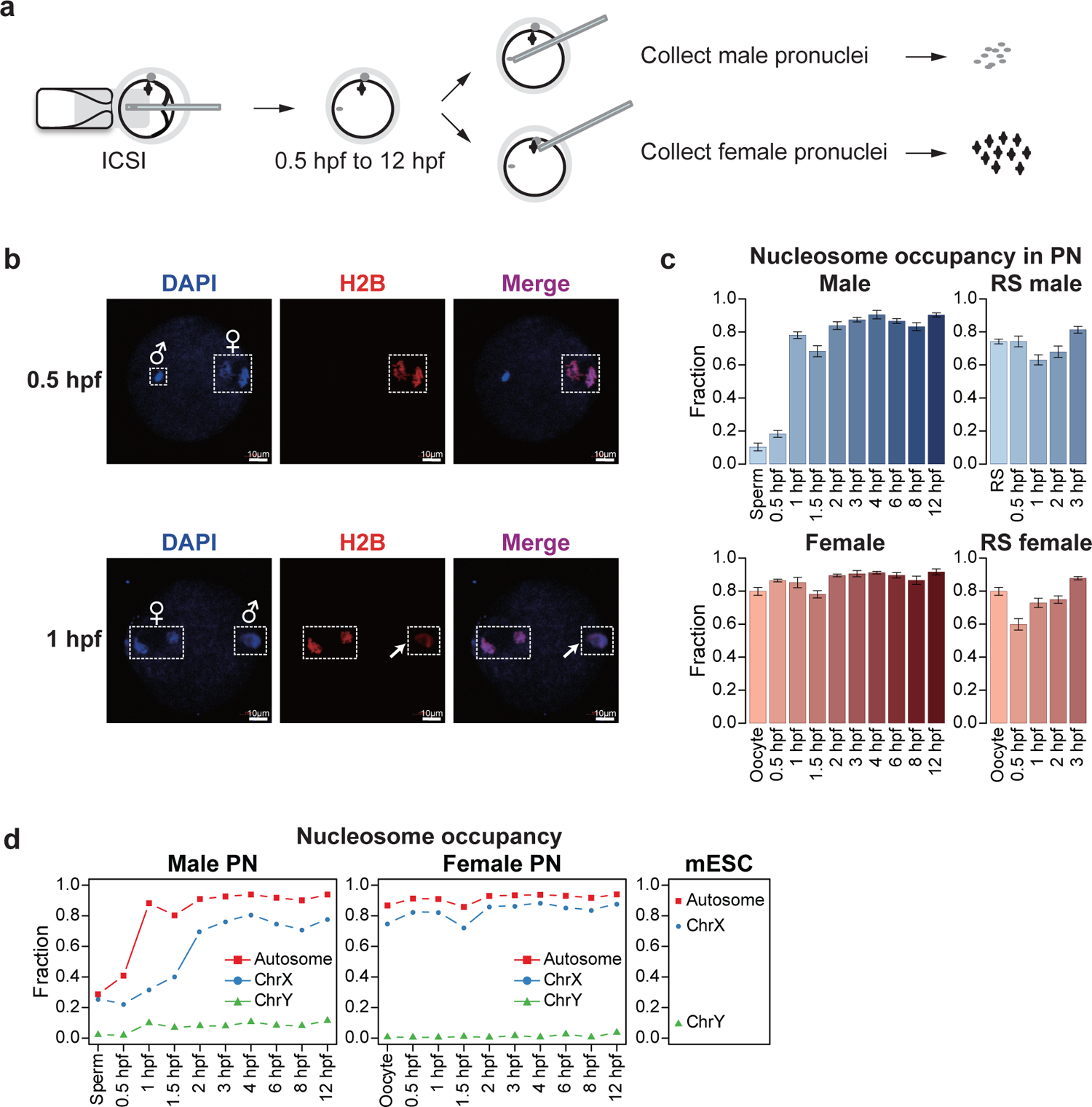
Nucleosome occupancy is quickly established in mouse male pronuclei after fertilization. **a** Schematic showing the collection of pronucleus samples for ULI-MNase-seq. **b** Confocal microscopy images of H2B-RFP mRNA-injected embryos shortly after fertilization. Newly incorporated H2B was present in the male PN (arrowhead) as early as 1hpf. **c** Bar plots showing the fraction of nucleosome-occupied 1-kb bins in each PN sample. Error bars represent ±1.96*SD. **d** Line charts showing the fraction of nucleosome-occupied 1-kb bins in sex chromosomes and autosomes of each PN and ESC samples. Chr, chromosome.

### Nucleosome occupancy is quickly established in the male pronuclei after fertilization

To characterize the chromatin remodeling process, NEPTUNE first identified nucleosome-occupied regions using a consecutive window approach; the resolution was set to 1 kb due to the sparseness of nucleosomes in PN samples (see Methods). Genome-wide nucleosome occupancy in oocytes and sperm was quite different from that of the PN samples, even in 0.5-hpf male PN and female PN (Fig. S3h). We next evaluated the nucleosome positioning dynamics in gametes and PN samples. In line with the previous study^22^, only 10-20% of the genome was occupied by nucleosomes in sperm and 0.5-hpf male PN, but this proportion dramatically increased to nearly 80% in 1-hpf male PN, indicating that nucleosomes were globally deposited into the paternal genome quickly at approximately 1 hpf (Fig. 1c). Importantly, this rapid *de novo* establishment process did not occur in female PN or parental PN from ROSI embryos (Fig. 1c), suggesting that this global nucleosome establishment was corresponding to the protamine-to-histone transition in the paternal genome after fertilization. In addition, we compared the genome-wide distribution of newly established nucleosomes for each stage. Consistently, the newly gained nucleosomes in PN stages were quite different from the nucleosomes in gametes. Retained nucleosomes in sperm were slightly enriched in promoters and short interspersed nuclear elements (SINEs) and telomeres, but the PN-established nucleosomes were more enriched in long interspersed nuclear elements (LINEs) (Fig. S4a). These results indicate that genome-wide chromatin remodeling occurs in both the paternal and maternal genomes after fertilization.

To investigate the function of nucleosome-occupied regions, we identified nucleosome-occupied promoters in sperm and promoters with newly established nucleosomes in 1-hpf and 6-hpf male PN (Fig. S4b). Gene ontology (GO) analysis showed that most genes with sperm-retained nucleosomes were associated with developmental process and cell differentiation (Fig. S4c), consistent with the discovery that sperm-retained histone modifications are enriched in promoters of developmental genes^23^. Genes obtaining nucleosome occupancy at 1 hpf in male PN were closely related to metabolic processes that are essential for cell replication and early embryonic development. Nucleosomes established at 6 hpf in male PN were enriched in genes involved in later organismal development such as sensory perception (Fig. S4c), which were rarely expressed at PN stages, indicating that the global nucleosome assembly occurred even at silent regions.

We next sought to investigate the difference of nucleosome establishment among different chromosomes. The percentage of nucleosome-occupied regions was lower in sex chromosomes than that in autosomes (Fig. 1d). Interestingly, the global establishment of nucleosome occupancy was also delayed on X chromosomes in the paternal genome, which occurred at 2 hpf. An earlier study suggested that oocyte TH2A/B variants were enriched in zygotes, especially in X chromosomes, which contributed to the activation of the paternal genome by inducing an open chromatin structure^24^. We hypothesized that the delay of the nucleosome occupancy in X chromosomes might result from the assembly of TH2A/B variants. We first validated the nucleosome occupancy on mESC-identified TH2A/TH2B peaks in all samples and found a higher MNase digestion sensitivity around the peak centers in sperm and the earlier stages of male PN, especially in X chromosomes (Fig. S4d). These results suggested a highly dynamic nucleosome assembly process at TH2A/TH2B peak regions. Moreover, the correlation between nucleosome occupancy and TH2A/B signal in X chromosomes appeared to be relatively high in 1-hpf male PN, but became negative in later male PN stages and all female PN stages (Fig. S4e), which also indicated the incorporation of TH2A/B variants in the paternal genome at earlier stages and a switch from TH2A/B to canonical nucleosome subunits at later male PN stages. Taken together, our results suggest that the acquisition of nucleosome occupancy in male PN is globally rapid and could be controlled by different histone variants.

### Distinct remodeling dynamics of NDRs in the paternal and maternal genomes

NDRs are usually highly accessible and corresponding to DNase I hypersensitive sites (DHSs) in the eukaryote genome, and they are typically located at regulatory regions including promoters, enhancers, and the origin of DNA replication^10^. The characteristic NDRs around TSSs provide binding hubs for transcription complex and are closely related to the regulation of gene expression^9^. In a recently published study in mice, the chromatin accessibility around TSSs was found to be increased greatly from gametes to zygotes, and nucleosome phasing was strongly positioned downstream of the +1 nucleosomes at the 2-cell stage^4^. However, when and how these proximal NDRs are established after fertilization remain to be unclear. We used NEPTUNE to generate profiles of average nucleosome density (see Methods) around TSSs for each sample (Fig. 2a). To our surprise, the dynamics of nucleosome positioning showed remarkable differences in maternal and paternal genomes. In male PN, the proximal NDR pattern appeared as early as 1.5 hpf, which was then enhanced and became more evident after 6 hpf (Fig. 2a). We observed a similar trend on the nucleosome phasing in male PN, in which the phasing periodicity was lost before 1 hpf and gradually rebuilt from 1.5 hpf (Fig. 2b-c and Fig. S5a). For the maternal genome, both the proximal NDRs and the nucleosome phasing were established at 3 hpf and became more obvious later at 6 hpf, and the nucleosome profiles became similar in the parental pronuclei after 6 hpf (Fig. 2a-c and Fig. S5a). These observations suggested that the nucleosome depletion pattern at promoters was generated earlier in the paternal genome than that in the maternal genome.

**Fig. 2.**
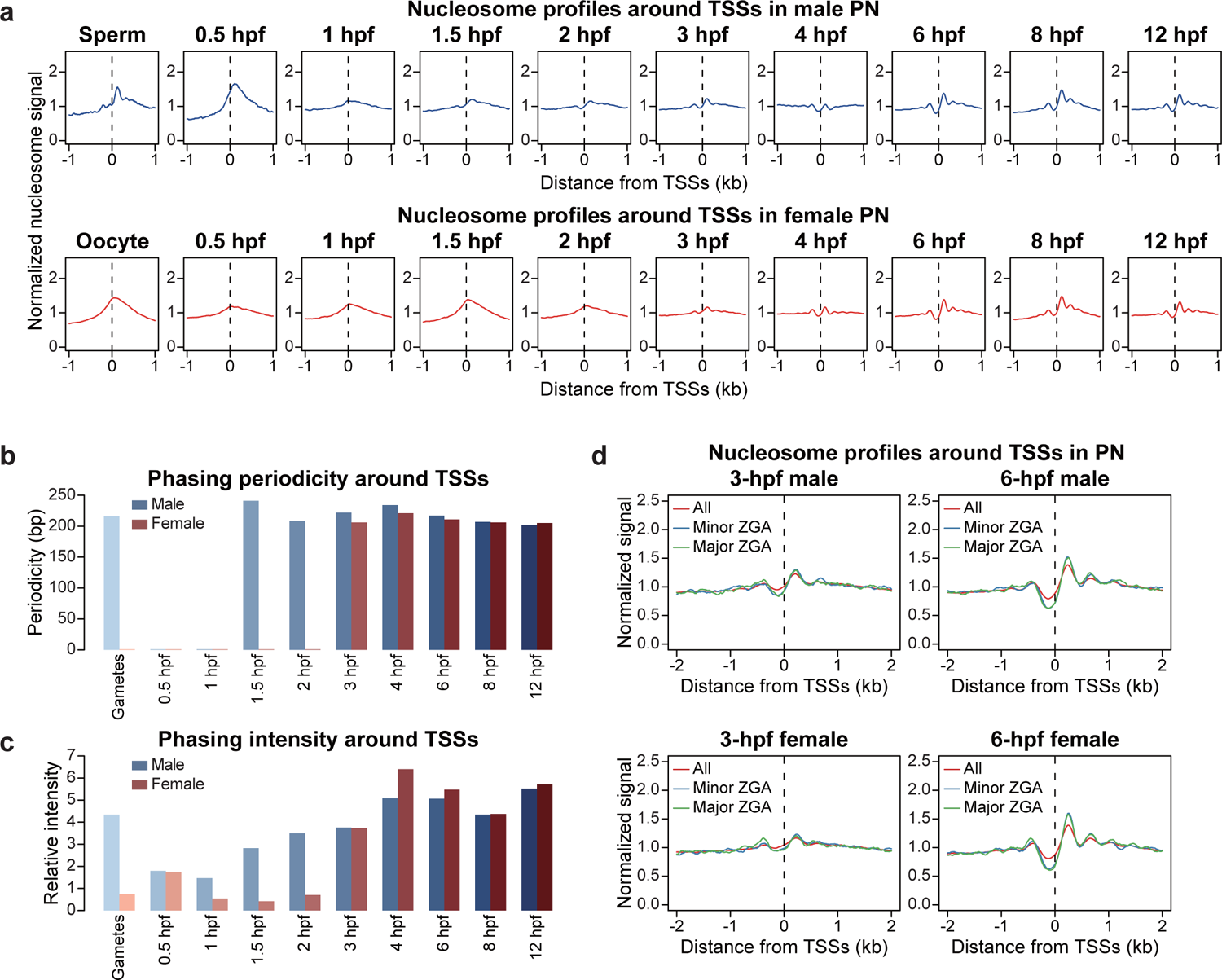
Asynchronous establishment of canonical nucleosome positioning in mouse parental pronuclei. **a** Nucleosome profiles around TSSs of Refseq genes at each PN stage. **b and c** Bar plots showing the nucleosome phasing periodicity (b; illustrated in Fig. S5a) or phasing intensity (c; calculated as the spectral intensity corresponding to the periodicity after fast Fourier transform) around TSSs of Refseq genes at each PN stage. **d** Nucleosome profiles around TSSs of all Refseq genes and ZGA genes in 3-hpf and 6-hpf parental PN.

As the nucleosome depletion pattern in parental genomes becomes general comparable for promoter regions, we asked whether this is true for other genomic loci, such as imprinted control regions (ICRs) and imprinted genes. The nucleosome profiles and NDRs are generally comparable between paternal and maternal genomes on ICRs (Fig. S5b). To quantify the nucleosome positioning dynamics, we calculated nucleosome depletion scores (NDR scores) and phasing scores (POS scores) for different gene sets, which evaluated the depth of NDRs and the periodicity of well-phased nucleosome arrays, respectively (see Methods). Interestingly, we observed an increase in NDR scores at 2 hpf for maternally imprinted genes especially in the maternal genome, and a decrease in NDR scores for paternally imprinted genes especially in the maternal genome (Fig. S5c), indicating that the imprinting control TFs such as CTCF might access the genome and initiate chromatin loops at this time. We also examined the formation of distal NDRs in enhancer regions. Considering that the enhancers in mouse pre-implantation embryos were established at relatively late stages, we identified these regions using ATAC-seq data from late 2-cell and inner cell mass (ICM) samples^25^. No significant NDRs were observed on late 2-cell-defined enhancers in either male or female PN (Fig. S5d), indicating that the chromatin remodeling on ZGA-related enhancers might be transient and occur after 12 hpf. Surprisingly, NDRs near the centers of ICM-defined enhancers were established as early as 6 hpf in parental PN (Fig. S5e), which was much earlier than the ICM stage when these functional enhancers were identified, suggesting that the pioneer factors regulating cell fates might start binding to the chromatin at as early as the PN stages. Unlike the distinct features of proximal NDRs, the dynamics of distal NDRs in the paternal and maternal genomes were much more similar.

We then analyzed whether the formation of NDRs was connected with gene expression. In mice, previous studies reported that the first wave of ZGA (designated the minor ZGA) began during the S to G2 phase at the 1-cell stage, and the second wave of ZGA (designated the major ZGA) occurred during the G1 phase at the mid-to-late 2-cell stage^7, 26^. We thus defined the significantly upregulated genes in zygotes compared to oocytes as minor ZGA genes, and upregulated genes in 2-cell-stage embryos compared to zygotes as major ZGA genes (see Methods). The NDR pattern was more obvious on promoters of both minor and major ZGA genes compared to all genes at 6 hpf when the minor ZGA begins (Fig. 2d). Strikingly, the NDR and nucleosome phasing patterns on ZGA genes were already more obvious at 3 hpf (Fig. 2d) when the genome should be quiescent and no transcription occurs, suggesting a priming effect of chromatin remodeling. Consistent with the previous observations, ZGA genes showed higher NDR scores and POS scores than average after 3 hpf in both male and female PN, and this difference became more significant at 6 hpf (Fig. S5f-g). These results suggest that the nucleosome positioning on promoters of ZGA genes is more strongly remodeled than other genes, which occurs before the start of transcription activation and might be important for ZGA initiation.

### GC content is a major determinant of nucleosome occupancy at pronucleus stages

Since the genome in male PN is quickly occupied by nucleosomes at 1 hpf, and the nucleosome positioning pattern also showed a remarkable difference between PN and gamete samples, we next explored the driving force responsible for rapid nucleosome occupation and remodeling. Multiple factors, including intrinsic sequence features, DNA and histone modifications, as well as active processes such as DNA replication, transcription, and activities of ATP-dependent chromatin remodelers, were found to impact nucleosome positioning^10, 27^. We first analyzed the sequence features of newly established nucleosomes at each stage and found that nucleosomes established earlier tended to have higher GC content (Fig. 3a), suggesting that nucleosomes preferred to occupy regions with high GC content, which was consistent with their intrinsic DNA sequence preference^28^. This trend was significant in the paternal genome, but not in the maternal genome or ROSI embryos, possibly due to the absence of the dramatic *de novo* nucleosome occupation in female PN or ROSI embryos (Fig. 3a and Fig. S6a-b). We next asked whether the chromatin state and histone modifications in sperm could impact the early establishment of nucleosome occupancy after fertilization. We calculated the partial correlation between normalized nucleosome occupancy and chromatin accessibility as well as DNA methylation state in early male PN. However, neither of them showed a strong correlation as GC content (Fig. S6c). These results suggest that the GC content, rather than the chromatin state of sperm, is the major determinant of nucleosome occupancy in early male PN. Next, we evaluated whether a higher GC content was also required for the establishment of promoter NDRs after fertilization. However, the correlation between newly established promoter NDRs and GC content was pretty low (Fig. S6d), suggesting that the NDR establishment on promoters was not mainly determined by intrinsic DNA sequence features. Additionally, neither the chromatin accessibility nor the DNA methylation level was correlated with the formation of promoter NDRs (Fig. S6e). The above analyses suggest that the intrinsic DNA sequence features might be essential for the nucleosome establishment, but have little effect on the nucleosome remodeling.

**Fig. 3.**
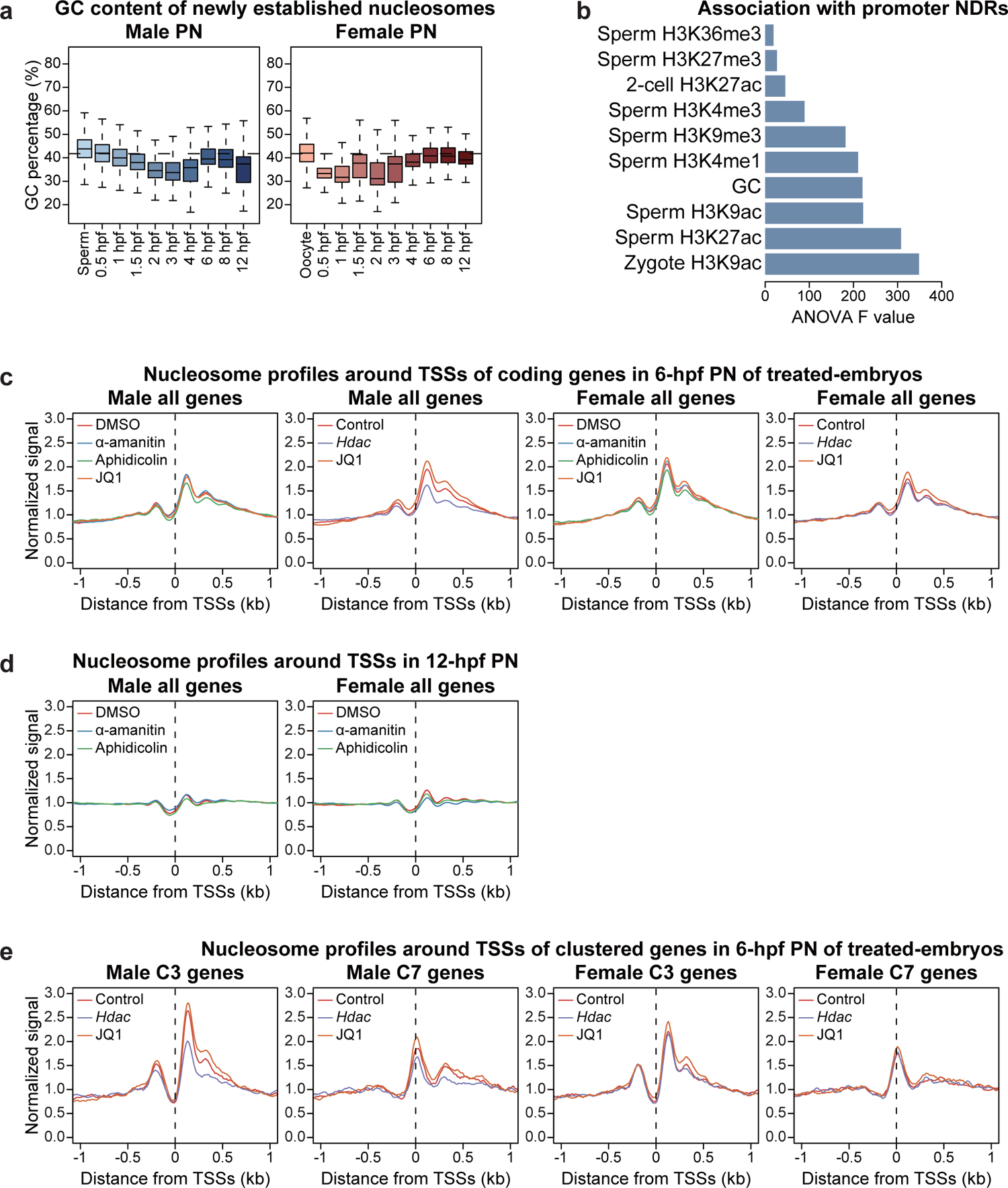
GC content and the histone acetylation level influence nucleosome establishment and repositioning respectively in mouse pronuclei. **a** Boxplots showing the GC content of newly established nucleosome regions at each PN stage. Dashed lines represent the average GC content in genome. b Bar plot showing the association of different histone modifications and GC content with promoter NDR scores. c, d and e Nucleosome profiles around TSSs of all Refseq genes at 6 hpf (c) and 12 hpf (d), or genes of indicated promoter NDR clusters at 6 hpf (e; defined in Fig. S6f) in parental PN from groups under different treatment. JQ1-B1: JQ1 batch1; JQ1-B2: JQ1 batch2. DMSO: Dimethyl sulfoxide; Control: Water-injected.

### Histone acetylation influences the establishment of NDRs in male PN

Next, we investigated the potential determinants of nucleosome repositioning after fertilization. We first characterized all genes based on NDR scores of promoters using k-means clustering, which revealed 7 clusters with differential NDR dynamics in male PN (referred to as C1-C7; Fig. S6f). The NDR scores of C1 and C5 were stable since sperm and inherited in latter stages; C2, C3, and C4 showed *de novo* establishment of NDRs at 1.5 hpf, 1 hpf, and 0.5 hpf, respectively; C6 and C7 showed weak NDRs in general. ANOVA analysis indicated that sperm- or zygote-identified histone acetylation was highly associated with NDR establishment (Fig. 3b), and clusters with high NDR scores (C3 and C4) also showed the highest level of H3K9ac and H3K27ac in both sperm and zygotes (Fig. S6g). These results suggest that histone acetylation might influence the establishment of NDRs at PN stages.

It was recently reported that in mouse embryos, blocking the elongation of RNA Pol II-mediated transcription by α-amanitin drastically compromised the openness of wider proximal NDRs^4^. Therefore, we treated embryos with α-amanitin or aphidicolin after ICSI to block transcription or DNA replication, respectively, considering that these two processes occur at PN stages^29^. Also, to evaluate the potential role of histone acetylation in the establishment of NDRs, we injected the mRNA of histone deacetylase gene *Hdac1* and *Hdac2* into MII oocytes and then performed ICSI. In addition, we used JQ1 to disrupt the binding of bromodomain proteins to acetyl-lysines^30, 31^. Inhibition of transcription or DNA replication in corresponding groups was confirmed by EU or EdU staining at 12 hpf, respectively (Fig. S6h-j). We then collected parental PN at 6 hpf from embryos under different treatments and performed ULI-MNase-seq with 2 or 3 biological replicates (Fig. S6k-l). We also collected α-amanitin or aphidicolin-treated samples at 12 hpf, when the transcription and DNA replication should have occurred in control samples (Fig. S6m). All of the above treatments had no significant influence on the global nucleosome occupancy compared to the control group, indicating a relatively stable protamine-to-nucleosome exchange (Fig. S7a-b). To our surprise, we observed little difference in nucleosome profiles of parental PN from α-amanitin- or aphidicolin-treated groups, and the promoter NDR scores were almost comparable to the control group at both 6 hpf and 12 hpf (Fig. 3c-d and Fig. S7c-d). The results were consistent on promoters of ZGA genes (Fig. S7e-f), although the transcription activity of ZGA genes were supposed to be blocked in the α-amanitin-treated group. In summary, these analyses suggest that the *de novo* establishment of NDRs is not mainly determined by transcription elongation or DNA replication activities in the first 12 hours after fertilization.

We then analyzed the nucleosome profiles of parental PN at 6 hpf from *Hdac* mRNA-injected or JQ1-treated groups. Interestingly, we found that after *Hdac* overexpression, the nucleosome positioning pattern on promoters was significantly changed with a decreased signal in +1 and +2 nucleosomes, and this disruption was more evident in male PN (Fig. 3c and Fig. S7g). Since the relative signal of the −1 and +1 nucleosome peaks is essential for defining the presence of a typical NDR, these observations suggest that *Hdac* overexpression might impede the formation of NDRs, which might further negatively regulate the transcription activity. The effects of JQ1 treatment in nucleosome profiles were not stable though, as JQ1-treated group showed higher promoter NDR scores compared to the DMSO-treated group in the first batch of data, but showed no significant difference compared with the control in the second batch of data (Fig. 3c and Fig. S7c). These results were further confirmed by nucleosome profiles on promoters of ZGA genes (Fig. S7e, g). In addition, nucleosome profiles of the JQ1-treated group highly resembled the control in both batches (Fig. 3c). *Hdac* overexpression causes a global reduction in histone acetylation and may lead to an altered chromatin state which is required for NDR formation, whereas JQ1 only inhibits Brd4-mediated recruitment of the preinitiation complex (PIC) on acetylated TSSs. Therefore, JQ1 might cause a weaker effect on the chromatin state compared to the genome-wide histone acetylation loss. We further generated the nucleosome profiles on promoters showing *de novo* NDR establishment in male PN after fertilization (C3), and compared them with profiles of promoters with weak NDRs throughout the PN stages (C7). Reassuringly, the signal of +1 nucleosomes on C3 promoters was significantly decreased in male PN upon *Hdac* overexpression, but C7 promoters did not share this change, and the nucleosome signal remained relatively stable on both C3 and C7 promoters in female PN (Fig. 3e). Taken together, our analyses suggest that histone acetylation, but not DNA replication or transcription, potentially guides the establishment of NDRs during the nucleosome incorporation process in male PN.

### NDR establishment on motif regions reveals TF binding dynamics in zygotes

The binding of TFs triggers transcriptional activation of the targeted genes by recruiting chromatin remodelers and RNA polymerase to promoter or enhancer regions^9, 10, 32, 33^. Although a subset of TFs can directly bind to nucleosomal DNA, many TFs have to compete with histones for binding to the motifs and creating open chromatin regions^9, 11^. In addition, previous studies suggested that NDRs and well-positioned nucleosomes were located around the TF binding sites, which were found to be correlated with the transcription activity of the targeted genes^12^. During the first cell cycle after fertilization, nucleosome remodeling permits the access of TFs to DNA, which could be important for the subsequent zygotic genome activation. Although several studies used DNase-seq or ATAC-seq to identify open regions in chromatin at zygote and 2-cell stages^3, 25^, it is still not feasible to make conclusions about the roles of TFs in nucleosome repositioning due to the insufficient input materials and limited sensitivity of these methods.

Our nucleosome profiling enabled us to examine the nucleosome repositioning process in TF motif regions shortly after fertilization, which might predict the binding status of specific TFs in the genome. In nucleosome profiles of mESC samples, strong NDRs and well-positioned nucleosome arrays around the motifs of CTCF have been observed (Fig. S1h), and these patterns were believed to be crucial for organizing chromatin structures in human embryos^34^. To our surprise, in male PN, typical NDRs on CTCF motif centers first appeared at 1.5 hpf (Fig. 4a), suggesting that CTCF might bind to its targets in the paternal genome at as early as 1.5 hpf. However, this process was delayed in the female PN which occurred at around 3 hpf (Fig. 4a), similar to the distinct formation of proximal NDRs around TSSs (Fig. 2a). Therefore, the NDR dynamics might suggest the potential binding of TFs on their motif sites. We then extended the analysis and calculated the NDR scores on motif centers of 122 TFs which were expressed at the zygote stage^35^. According to the dynamics of NDR scores, we divided these TFs into three clusters using k-means clustering (Fig. 4b). For cluster-1 TFs, the NDR scores on their motifs remained low in all stages, indicating that the binding sites of these TFs were exclusively covered by nucleosomes (Fig. S8a). In contrast, motifs of cluster-2 TFs maintained high NDR scores, on which nucleosomes were depleted throughout the PN stages (Fig. S8b). We further found that the high or low abundance of nucleosomes on motifs of cluster-1 or cluster-2 TFs might be determined by the GC content of these motif sequences (Fig. S8c). Interestingly, for motifs of cluster-3 TFs including CTCF, although the GC content was as high as motifs of cluster-1 TFs (Fig. S8c), the NDR scores increased after fertilization in both the paternal and maternal genomes, with the paternal genome increased earlier (Fig. 4b), indicating that an active nucleosome repositioning process created NDRs at these TF binding sites. These results indicate that cluster-3 TFs might access the genome and be involved in chromatin remodeling or even transcription activation at the zygote stage.

**Fig. 4.**
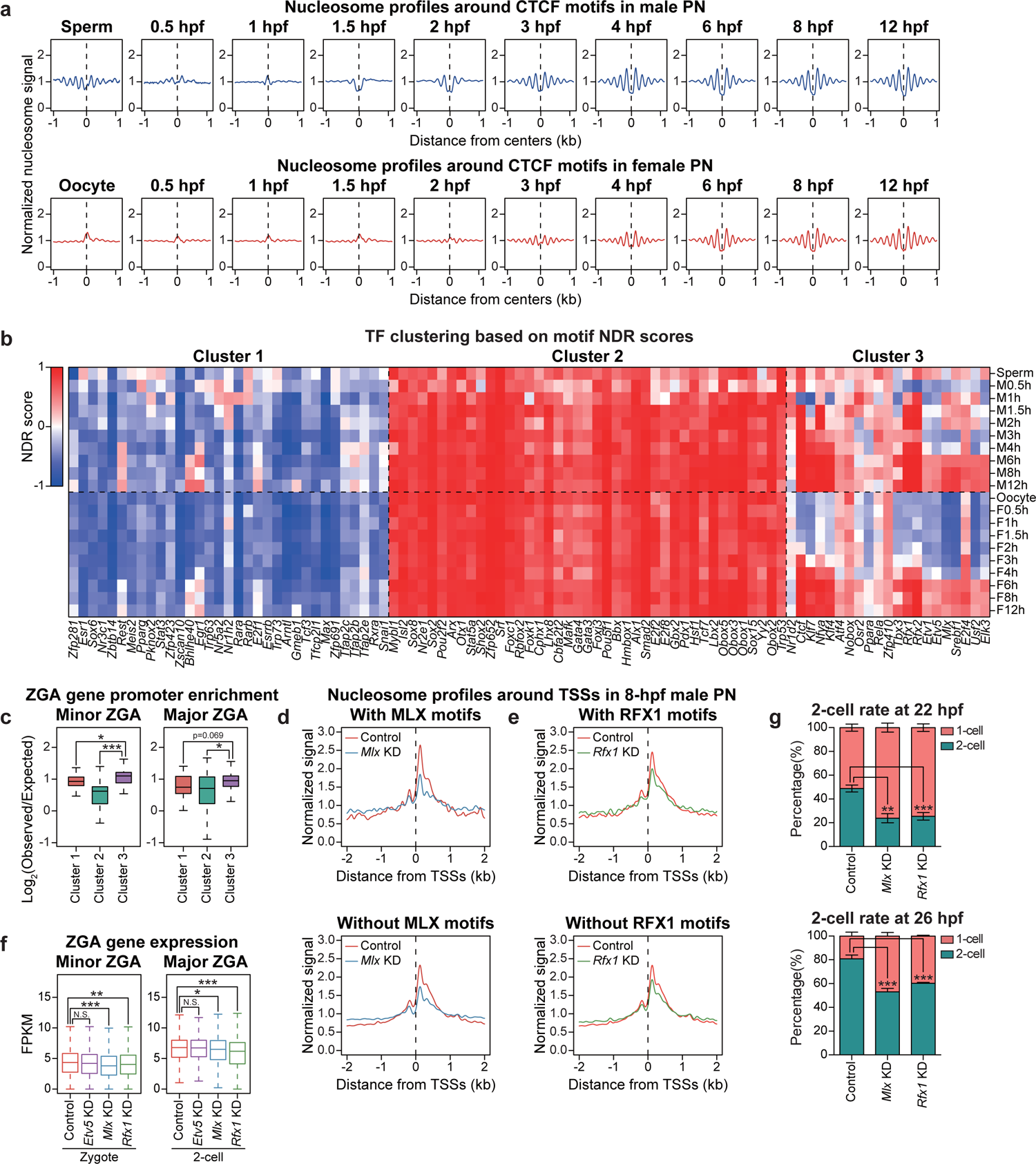
Dynamics of nucleosome organization on motif regions predict TF binding landscapes in mouse pronuclei. **a** Nucleosome profiles around CTCF motifs at each PN stage. **b** Heatmap showing the k-means clustering (k=3) of TFs based on NDR scores on motifs at each PN stage. M, male PN. F, female PN. h, hpf. **c** Boxplots showing the enrichment of ZGA gene promoters on motifs of TFs in different clusters defined in b (*** p < 0.001; * p < 0.05). **d and e** Nucleosome profiles around TSSs of different classes of genes in 8-hpf male PN from KD groups. Genes were classified according to whether motifs of MLX (d) or RFX1 (e) are present in promoters. **f** Boxplots showing the expression level of ZGA genes in KD embryos (*** p < 0.001; N.S. p > 0.05). **g** Stacked bar plots showing the percentage of 1-cell and 2-cell embryos in KD groups at indicated time points. n=3 biological replicates with approximately 50 embryos each. Error bars represent SD.

### MLX and RFX1 promote NDR establishment in zygotes and ZGA

We speculated that cluster-3 TFs, which seemed to access chromatin at as early as PN stages, were related to the regulation of ZGA. To validate this hypothesis, we first calculated the enrichment of ZGA-associated promoters on TF motifs for each cluster. As expected, compared to cluster-1 and cluster-2 TFs, promoters of minor and major ZGA-related genes were significantly more enriched on motifs of cluster-3 TFs (Fig. 4c). We further identified 13 candidate TFs from cluster 3 whose motifs showed significant enrichment of ZGA-associated promoters (Fig. S8d), including NFYA which is proved to contribute to the ZGA process and the formation of DHSs at the 2-cell stage^3^. Therefore, the predicted candidates whose binding sites showed similar nucleosome positioning dynamics with NFYA might also play a role in regulating ZGA. To further narrow down the candidates, we compared the expression pattern of these TFs during mouse embryonic development (Fig. S8e). As an important regulator in ZGA, *Nfya* was highly expressed in oocytes and zygotes, indicating that NFYA was maternally stored. However, only 7 of the 13 candidates including *Nfya* showed a relatively high maternal storage (*Etv1*, *Nfya*, *Usf2*, *Klf7*, *Srebf1*, *Mlx*, *Rfx1*; Fig. S8e). Among these predicted TFs, we selected a minor ZGA-related TF MLX and a major ZGA-related TF RFX1 for further verification. MLX is a glucose-sensing transcription factor which translocates from the cytoplasm to the nucleus and binds to specific DNA motifs in response to the glucose stimulus, leading to an increased histone H4 acetylation level at target promoters and activation of corresponding genes^36^. RFX1 regulates various kinds of genes including ribosomal genes, tissue-specific genes and cellular communication-associated genes^37–40^. Importantly, homozygous knockout of *Rfx1* leads to early embryonic lethality before implantation^41^, suggesting an indispensable role of *Rfx1* in early development. As shown, the nucleosome depletion pattern appeared at approximately 6 hpf around the binding motifs of MLX and RFX1, and these two factors were both maternally expressed (Fig. S8e and S9a-b). Here, we also used ETV5 as a negative control, the motif of which showed a high enrichment of ZGA-associated promoters but was barely expressed in oocytes or zygotes (Fig. S8d-e).

To reduce the impact of maternal proteins, we injected siRNAs targeting random sequences or TFs into GV-stage oocytes and performed ICSI after *in vitro* maturation. We first applied ULI-MNase-seq to parental PN of knockdown (KD) embryos at 8 hpf (Fig. S9c) and evaluated the effect on NDR establishment. To reduce the noise generated by differences in embryo culturing and KD efficiency, we prepared 2-3 biological replicates for each KD group and averaged the nucleosome profiles for downstream analyses. Surprisingly, KD of *Mlx* and *Rfx1* altered the nucleosome profiles on promoters of male PN, but had little effect for female PN (Fig. 4d-e and Fig. S9d-e), which might be caused by the differential time course and/or protein participation of chromatin remodeling in parental PN. Moreover, promoters possessing motif sequences for MLX or RFX1 showed greater changes on the nucleosome profiles in male PN, with more severely decreased +1 and −1 nucleosomes (Fig. 4d-e and Fig. S9f). We further evaluated the nucleosome profiles on promoters of minor and major ZGA genes, and observed a decrease in +1 and −1 nucleosomes compared to other genes in male PN of KD embryos (Fig. S9g-h). In summary, the decreased signal of +1 and −1 nucleosomes in KD embryos suggest a reduction in NDR formation and transcription activity, indicating that in male PN, MLX and RFX1 might be responsible for creating NDRs on promoters of ZGA genes. We next asked whether the reduction in NDR formation at promoters of male PN would affect the ZGA in mice. We then collected control, *Etv5*, *Mlx*, or *Rfx1* KD embryos at late 1-cell (16 hpf) and late 2-cell (40 hpf) stages for RNA-seq. We first confirmed that the expression level of these factors was significantly reduced and the replicates were highly consistent (Fig. S10a-b). Interestingly, little transcriptome difference was observed when *Etv5* was silenced, but significantly more genes were differentially expressed in *Mlx* KD zygotes, and a large number of genes were downregulated in *Rfx1* KD 2-cell embryos (Fig. S10c-d). Notably, minor and major ZGA genes were significantly downregulated after *Mlx* or *Rfx1* silencing, respectively (Fig. 4f). Functional analyses also suggested that downregulated genes in *Mlx* or *Rfx1* KD embryos were highly enriched in ZGA-associated processes and genes with MLX or RFX1 motifs (Fig. S10e-f). Consistently, although the overall proportion of embryos able to reach the 2-cell stage in the KD groups was comparable with the control (Fig. S10g), silencing of *Mlx* or *Rfx1* significantly prolonged the 1-cell stage (Fig. 4g), indicating that the ZGA process was delayed in the KD embryos. Taken together, our data demonstrate that the predicted factors MLX and RFX1 are possibly required for the timely completion of ZGA through regulating the establishment of promoter NDRs on corresponding genes.

## Discussion

In this study, we analyzed the dynamic changes in genome-wide nucleosome occupancy and positioning in mouse embryos during the first 12 hours after fertilization. We traced the differential changes in the parental genome and started our observations as early as 30 min after fertilization by ICSI. We assessed the detailed pattern of NDR rebuilding on promoters and nucleosome positioning dynamics on TF binding motifs, which uncovered novel molecular regulation mechanisms for the ZGA process in mice.

Multiple epigenetic landscapes have been linked to transcription activation and subsequent regulations in mammalian embryos, including histone modifications, DNA modifications, chromatin accessibility, and high-dimensional structures^7, 42^. However, the fundamental issues, hierarchy, and connected factors of the maternal-to-zygotic transition remain unclear due to the difficulties in setting up proper experimental strategies and analyses with high sensitivity. To tackle this problem, we profiled the nucleosome positioning at the very early stages after fertilization to study the initiation process of chromatin reorganization. We found that the NDR establishment on promoters appeared at as early as 1.5 hpf in the paternal genome and 3 hpf in the maternal genome, both of which were earlier than the transcription activation at the PN3 stage (6 hpf); further, the profiles of promoter NDRs were equalized in the parental genomes after 6 hpf. Interestingly, this discrepancy in the creation of promoter NDRs was very similar to the differential H4 hyperacetylation dynamics and transcription activities in the parental pronuclei revealed by immunofluorescent staining^18^. Meanwhile, our analyses suggest that histone acetylation, but not DNA replication or RNA Pol II elongation, might be crucial for the re-establishment of promoter NDRs and +1 nucleosomes upon fertilization. Reducing the histone acetylation level by *Hdac* overexpression attenuated the formation of promoter NDRs only in male PN, which might be caused by the cascade effect on chromatin state alterations and reduced binding of pioneer transcription factors, as the male genome is opened earlier than the female genome. Blocking the recognition of histone acetylation by JQ1 showed a weaker and unstable influence on NDR formation and +1 nucleosomes, which also suggested that the Pol II recruitment or transcription initiation might not be required for creating NDRs at the PN stages. On the other hand, generation of promoter NDRs might be crucial for initiating the ZGA process, and further investigations are required to clarify factors responsible for the subsequent remodeling on promoter NDRs in both paternal and maternal genomes, which possibly relies on the function of ATP-dependent chromatin remodelers such as the SWI-SNF complex.

It is generally believed that in the ZGA model, some pioneer TFs are able to initiate the transcription activation, but how to identify these pioneer factors for ZGA remains a long-standing question^7, 43^. Interestingly, in the somatic cell reprogramming induced by TFs, pioneer factors also have the ability to access partial motifs on nucleosomal DNA and gradually change the chromatin status from silent to open^44^. In our study, we provided a new strategy, NEPTUNE, which includes functions for predicting pioneer factors during the developmental process *in vivo*. NEPTUNE first identified the dynamics of nucleosome positioning on TF binding motifs and screened the close-to-open transition to predict pioneer factors that might bind the nucleosome-occupied chromatin and gradually open the genome with the help of other chromatin remodelers. Based on our data from PN-stage samples, NEPTUNE identified dozens of TFs whose motif regions showed obvious changes in the chromatin state, including the previously reported factor NFYA, as well as the novel regulator MLX and RFX1.

Blocking the function of MLX and RFX1 leads to attenuated NDR formation and failed activation of certain ZGA-associated genes through either direct or indirect mechanisms. MLX was found to activated myokines by increasing the histone acetylation level^36^, and this glucose-sensing factor could balance metabolism to suppress apoptosis and promote cell survival^36^. RFX1 could also retain the cell viability under stress through activating cellular communication network factors^40^. The ZGA process also induces many metabolism-associated genes, providing the possibility that MLX and/or RFX1 contribute to these transcriptional activation events. Although MLX and RFX1 both regulate metabolic genes, their motifs and target genes are highly different (Fig. S10h-i). In addition, both RFX1 and NFYA regulate a subset of major ZGA genes, also with few overlaps between each other (Fig. S10j). These analyses suggest that the ZGA regulation is highly complex and may require the involvement of multiple TFs. We are working on the construction of oocyte-specific knockout mouse models for MLX and RFX1 to systematically investigate their roles in facilitating ZGA at the early stages. Collectively, our data provide rich resources for the study of the mechanisms involved in ZGA, and the NEPTUNE method may also be helpful for exploring the epigenetic regulation in other developmental events after fertilization.

## Materials and Methods

### Animals and collection of mouse embryos

Specific-pathogen-free (SPF) mice were housed in the animal facility at Tongji University, Shanghai, China, and they were fed a standard diet. The temperature and light were strictly controlled (24°C; 12 hours light and 12 hours dark). All animal experiments were performed in accordance with the University of Health Guide for the Care and Use of Laboratory Animals and were approved by the Biological Research Ethics Committee of Tongji University. B6D2F1 female mice (8–10 weeks old) were superovulated by injection with 7 IU of pregnant mare serum gonadotropin (PMSG), which was followed by injection with 5 IU of human chorionic gonadotropin (hCG) (San-Sheng Pharmaceutical Co., Ltd) 48 hours later. MII oocytes were collected from the oviducts of the superovulated female mice.

### Mouse sperm extraction and ICSI

Both cauda epididymides were collected from each C57BL/6 male mouse and then were punctured by needles. The semen was then squeezed out and placed into a 1.5 ml Eppendorf tube containing 500 μL of warm HEPES-buffered CZB (HCZB) medium; the sample was then incubated at 37°C for 10–15 min to allow sperm to swim out. ICSI was then performed on the stage of an Olympus inverted microscope equipped with a Narishige micromanipulator. MII oocytes were placed in a drop of HCZB medium, and a single sperm head was injected into each MII oocyte with the aid of a piezo-driven micromanipulator. Embryos were then cultured in G-1 PLUS medium (Vitrolife) after fertilization before harvesting.

### H2B-RFP overexpression followed by immunostaining in embryos

H2B cDNA fused with the sequence of RFP was cloned into a T7-driven vector, and H2B-RFP mRNA was synthesized with the mMESSAGE mMACHINE T7 Transcription kit (Invitrogen, AM1344) following the manufacturer’s instructions. The concentration of the injected mRNA was found to be optimal at 100 ng/μL. MII oocytes were injected with approximately 10 pl of the diluted mRNA using a piezo-driven micromanipulator. After the injection, the oocytes were cultured for 2 hours to allow recovery and H2B-RFP expression before fertilization.

At specific time points after ICSI, fertilized embryos were fixed with 4% paraformaldehyde in PBS for 1 hour at room temperature (RT). The fixed embryos were then washed in 0.5% BSA in PBS and treated with 0.2% Triton X-100 for 20 min at RT for permeabilization. The nuclei were stained with DAPI for 10 min at RT before the embryos were mounted on a glass slide in anti-bleaching solution. Fluorescence was detected under a laser-scanning confocal microscope (Zeiss, LSM880).

### Isolation of parental pronuclei after fertilization

At 0.5 hpf, 1 hpf, 1.5 hpf, 2 hpf, 3 hpf and 4 hpf, the embryos were placed into HCZB medium containing Hoechst 33342 dye to make the pronuclei visible. At time points later than 4 hpf, the pronuclei became visible without staining. Zona pellucidae were punctured with a piezo-drill micromanipulator, and the pronuclei were isolated from the embryos. The parental pronuclei were distinguished by their sizes and distances from the second polar bodies. Isolated pronuclei were washed with 0.5% BSA in PBS before they were placed into the lysis buffer for low-input MNase-seq.

### Ultra-low-input MNase-seq

10-15 pronuclei per replicate were isolated and washed before they were placed into 0.7 μL of lysis buffer (10 mM Tris-HCl pH 8.5, 5 mM MgCl_2_, 0.6% NP-40) for individual reactions. Then, 2.5 μL of MNase master mix (MNase Buffer, 0.125 U/μL MNase (NEB, M0247S), 2 mM DTT, and 5% PEG 6000) was added into each tube, and the reaction was incubated at 25°C for 10 min for chromatin fragmentation. The reaction was stopped by the addition of 0.32 μL of 100 mM EDTA, and then 0.32 μL of 2% Triton X-100 was added to the reaction to release the fragmented chromatin. Then, 0.2 μL of 20 mg/ml protease (Qiagen) was added, and the reaction was incubated at 50°C for 90 min for protein digestion followed by incubation at 75°C for 30 min for protease inactivation.

The sequencing libraries were prepared using the KAPA Hyper Prep kit for the Illumina platform (kk8504) following the manufacturer’s instructions. After standard procedures including end repair and A-tailing, adapter ligation, post-ligation cleanup and library amplification, the resulting products were subjected to a second round of PCR amplification with the same provided primers to generate sufficient DNA materials for high-throughput sequencing. Paired-end sequencing with 150-bp read length was performed on the HiSeq X Ten (Illumina) platform at Cloudhealth Medical Group Ltd.

### Treatment of α-amanitin, aphidicolin or JQ1 and *Hdac* overexpression

For drug-treated groups, embryos were placed to G-1 PLUS medium supplemented with 0.05% DMSO, 100 μM α-amanitin (Sigma-Aldrich, A2263), 3 μg/mL aphidicolin (Sigma-Aldrich, A4487) or 1 μM JQ1 (MedChem Express, HY-13030) respectively after ICSI-induced fertilization. For *Hdac* overexpression, the mRNA of *Hdac1* and *Hdac2* was synthesized as described above and mixed at a concentration of 500 ng/μL each, and the *Hdac* mRNA mixture was injected into MII oocytes before ICSI. Embryos injected with water served as the control group for *Hdac* overexpression. At 6 hpf, the parental PN were collected for low-input MNase-seq.

To verify that the transcription or DNA replication activities were successfully inhibited in each group, we transferred embryos into G-1 PLUS medium supplemented with corresponding drugs as well as 1 mM EU or 10 μM EdU at 8 hpf, and cultured these embryos for another 4 hours. At 12 hpf, we fixed the embryos with 4% paraformaldehyde in PBS and performed EU (Invitrogen, C10329) or EdU staining (Invitrogen, C10634) respectively following the manufacturer’s instructions.

### Knockdown by siRNA injection in GV-stage oocytes and *in vitro* maturation (IVM)

Two or three siRNAs were designed for each gene, and the sequences were listed as follows: control siRNA (siCtrl; UUCUCCGAACGUGUCACGUTT); siEtv5-1 (UACAUGAGAGGCGGGUAUUUC); siEtv5-2 (AGCUUGCCCUUUGAGUAUUAU); siEtv5-3 (GCGACCUUUGAUUGACAGAAA); siMlx-1 (CGGUGUCCUUCAUCAGUUGAA); siMlx-2 (GAAAGUGAACUAUGAGCAAAU); siRfx1-1 (AGAACACUGCACAGAUCAA); siRfx1-2 (ACUGUGACAAUGUGCUGUA); siRfx1-3 (UCAUGGUAAACCUGCAGUU). The siRNAs against each gene were mixed together and diluted at a total concentration of 20 μM. Ovaries were obtained from B6D2F1 female mice (8–10 weeks old) 48 hours after PMSG injection and were then transferred to M2 medium (Sigma-Aldrich, M7167) supplemented with 0.2 mM 3-isobutyl-1-methylxanthine (IBMX; Sigma-Aldrich, I5879). The ovarian follicles were punctured with a syringe needle, and GV-stage oocytes were collected using a narrow-bore glass pipette. The GV-stage oocytes were then injected with approximately 10 pl of siRNAs using a piezo-driven micromanipulator.

For IVM, the injected GV oocytes were washed thoroughly in IBMX-free αMEM (Sigma-Aldrich, M0446) and were then incubated for 16 hours in the maturation medium (5% FBS and 1.5 IU/ml hCG in αMEM). Oocytes presenting with a polar body were classified as MII, and ICSI was then performed to fertilize oocytes at the designated time. At 22 hpf and 26 hpf, the percentage of 2-cell embryos in each group was calculated as the early 2-cell rate. At 48 hpf, the percentage of embryos at 2-cell or 4-cell stages in each group was calculated as the overall 2-cell rate. At 16 hpf and 40 hpf, late 1-cell and late 2-cell embryos were harvested respectively for RNA-seq.

### RNA-seq library generation

RNA-seq libraries were prepared as previously described^45^. Briefly, harvested blastomeres were placed in lysis buffer containing 0.5% Triton X-100, free dNTPs and tailed oligo-dT oligonucleotides. Reverse transcription was performed with Superscript II (Invitrogen 18064014), and cDNA amplification was performed as described. The amplified cDNA was fragmented using a Covaris sonicator (Covaris S220) with conditions as follows: peak power 50, duty factor 20, cycles/burst 200, 2 min. The KAPA Hyper Prep kit (kk8504) was applied to generate sequencing libraries following the manufacturer’s instructions.

### NEPTUNE pipeline for analyzing ULI-MNase-seq data

We developed a computational pipeline NEPTUNE (i**N**tegrat**E**d **P**ipeline **T**o analyze **U**ltra-low-input **N**ucleosome s**E**quencing data) for integrated analysis of ULI-MNase-seq datasets. NEPTUNE consists of four major steps described as followed (Fig. S2), and is freely available at https://github.com/chenfeiwang/NEPTUNE.

### Step1: Data pre-processing

#### Data pre-processing

MNase-seq reads were aligned to the mouse genome build mm9 using the bwa (v 0.7.12) mem command^46^. Reads with MAPQ less than 10 were removed from downstream analyses. To create nucleosome profiles, we identified the centers of all paired-end reads and extended them to 146-bp lengths. For nucleosome profile visualization, the middle 74 bp were compiled using the “genomeCoverageBed” function of bedtools^47^; for nucleosome occupancy calculation, the whole fragment was piled up. To normalize the effect of sequencing depth, all nucleosome profiles were scaled to 500 million reads in total.

#### Quality control

NEPTUNE randomly sampled 1M pair-ended reads of each sample, and calculated the distance between paired ends as the read length to generate the length distribution plot. NEPTUNE also calculated the genome-wide nucleosome coverage by enumerating the 200 bp bins which were covered by the nucleosome signal. To examine the reproducibility of the MNase-seq libraries, we generated nucleosome profiles for all replicates and calculated the correlation of normalized nucleosome occupancy between biological replicates using promoter regions (defined as 2 kb upstream and downstream of TSSs) of Refseq genes. As the replicates were highly correlated with each other (Pearson’s correlation > 0.8, except for nucleosome profiles from sperm or 0.5-hpf male PN samples, for which the low correlation might be caused by random nucleosome presence in the genome), we pooled the biological replicates together for each stage. Finally, NEPTUNE generated the averaged nucleosome profiles around unique regions such as TSSs, enhancers and the top 10000 CTCF binding regions.

### Step 2: Nucleosome occupancy and positioning modeling

#### Definition of normalized nucleosome occupancy

To calculate the genome-wide nucleosome occupancy for mESCs and parental pronuclei, we first separated the genome into 1 kb consecutive bins. Although we normalized the total sequence depth to 500 million reads per sample, in some samples such as sperm, 0.5-hpf male PN as well as single-cell ESC samples, only 5% to 30% of the genomes were occupied by nucleosomes, leading to a higher background noise. To normalize the background noise, we took the genome-coverage fraction into consideration, and the relative nucleosome occupancy O was defined as

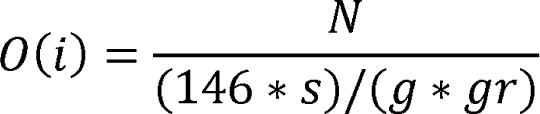

where N represents the number of normalized nucleosome fragments in this 1 kb region, s represents the normalized sequence depth (500 million reads), g represents the mouse genome size (2.7E9), and gr represents the fraction of genome occupied by nucleosomes, which is variable for different samples. The O(i) thus represents the number of observed nucleosome fragments versus the number of expected nucleosome fragments at the designated region. We calculated the relative occupancy, O, for each bin from ESC MNase-seq samples using different amounts of starting materials and estimated that bins with O(i) > 0.3 could represent nucleosome-occupied regions. We then used the same cut-off for parental PN samples to determine the nucleosome-occupied bins. Sperm-retained nucleosomes were defined as regions with O(i) >3 in sperm samples. Newly established nucleosome regions in each stage were defined as regions containing no nucleosome (O(i) <= 0.3) in any of the previous stages, but containing nucleosomes (O(i) > 0.3) at the present stage.

#### Nucleosome profiles around TSSs, ZGA genes and transcription factor motif regions

We generated the averaged nucleosome profiles around TSS regions and TF binding motifs using the SitePro function from CEAS^48^. TSS regions were profiled 2 kb upstream and downstream of the TSSs. TF binding regions were profiled 1 kb upstream and downstream of the motif centers, and only the top 10000 regions with significant motif scores were included in the profiles. Enhancer regions were defined using ATAC-seq peaks at the corresponding stages, excluding the peaks from promoter regions^25^.

#### Definition of nucleosome depletion score (NDR score), phasing score (POS score), and NDR loss ratio

To evaluate the nucleosome depletion and phasing status of TSS regions as well as TF binding motifs, we defined the NDR score and POS score.

The NDR score was defined as

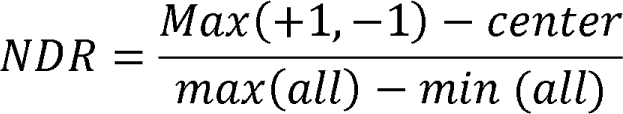

where +1 represents plus one nucleosome, which is the max of the normalized nucleosome profile from +50 bp to +250 bp of the TSS or motif center; −1 represents minus one nucleosome, which is the max of the normalized nucleosome profile from −250 bp to −50 bp of the TSS or motif center; the center represents the center region, which is defined as the mean of the normalized nucleosome profile from −50 bp to 50 bp; all represents all of the profiles, which represents −2 kb to +2 kb of the TSS or −1 kb to +1 kb of the motif center. The depletion score DS thus represents the depth of nucleosome depleted regions (NDR) versus all profiles, which should range from −1 to 1. If the NDR score is larger than 0, then there is a canonical nucleosome-depleted region at the center; if the NDR score is less than 0, then the center region is assuredly occupied by nucleosomes.

The phasing score is defined as

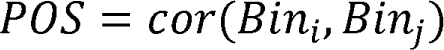

where bin i represents a bin from the TSS or the center of a motif to 1 kb downstream, with a 50 bp resolution; bin j represents a bin from the TSS or the center of a motif +10 bp to 1 kb downstream, with a 50 bp resolution. The correlation between these two bins, ranging from −1 to +1, represents the periodicity of the profile.

### Step 3: Perturbation evaluation

#### Generation of perturbated profiles

NEPTUNE generated the averaged nucleosome profiles around TSSs of all genes or ZGA-associated genes as described in Step 2. Users could also use a custom-defined gene list to generate the nucleosome profiles around TSSs. To normalized the influence of sequencing depth on nucleosome profiles and to compare profiles of different groups with each other, we divided the nucleosome signal at specific sites (calculated based on counts of MNase-seq reads) by the averaged signal intensity in the corresponding sample. For nucleosome profiles of TF KD groups which were generated using a small set of genes, we smoothed the nucleosome profiles using smooth.spline function in R for better visualization. Quantification of difference in NDR scores. NEPTUNE calculated the NDR scores of TSSs for differentially treated groups using the formula in Step 2. Significance between different groups was evaluated using the one-sided Wilcoxon rank-sum test.

### Step 4: Regulator screening

#### Clustering of transcription factors based on NDR dynamics at their binding motifs

To classify the functions of different TFs during the chromatin remodeling in mouse early embryogenesis, NEPTUNE calculated the nucleosome depletion scores for the TF motifs from the Cistrome database, including 335 curated motifs revealed by ChIP-seq data^35^. We only focused on the TFs expressed at the oocyte, zygote, early 2-cell and late 2-cell stages, and 122 TFs were left after setting the cut-off for FPKM as 1. For each TF, the corresponding binding sites were defined using the top 10000 sites with highest motif scores across the genome. We then calculated the NDR scores on these sites for all the parental PN stages and performed k-means clustering setting k = 3, which identified TFs with binding sites that 1) were always occupied by nucleosomes at the PN stages; 2) always had NDRs at the PN stages; 3) had a transition from nucleosome-occupied regions to NDRs, which might be resulted from the binding of corresponding TFs during embryogenesis.

#### ChIP-seq, ATAC-seq, RNA-seq data processing and normalization

Public sperm histone modification data from ChIP-seq and ATAC-seq experiments were used in the analysis^23^. ChIP-seq and ATAC-seq reads were aligned to the mouse genome build mm9 using the bwa (v 0.7.12) mem command^46^. Signal tracks for each sample were generated using the MACS2 (v2.0.10.20131216) pile-up function and were normalized to reads per million mapped reads (RPM)^49^. The RNA-seq reads from the knockdown (KD) experiments were mapped to the mm9 reference genome using STAR (v2.5.2b)^50^. Expression levels for all Refseq genes were quantified to fragments per kilobase million (FPKM) using stringtie (v1.3.6)^51^, and FPKM values of replicates were averaged.

#### Genomic enrichment analysis of nucleosome regions

The enrichment of nucleosome regions on genomic elements including promoters, high CpG density promoters (HCPs), intermediate CpG density promoters (ICPs), low CpG density promoters (LCPs), exons, introns, long interspersed nuclear elements (LINEs), short interspersed nuclear elements (SINEs), and long terminal repeats (LTRs) was calculated using observed probability versus expected probability. The observed probability was calculated using the lengths of nucleosome regions that covered the designated genomic elements versus the lengths of total nucleosome regions, and the expected probability was calculated using the total lengths of designated genomic regions versus the length of the whole genome.

#### Gene ontology (GO) analysis

Functional annotation analysis was performed using the MAGeCK-Flute package^52^. We only selected the Gene Ontology terms from biological processes to calculate the enrichment. P-values were calculated similar to the online tool of DAVID, which is based on a modified Fisher’s exact test.

#### Imprinting control regions and imprinted genes

We obtained 179 known imprinted genes (267 transcripts) from the geneimprint website (http://www.geneimprint.com) and previous publications^53, 54^. All transcripts were separated into maternally imprinted and paternally imprinted based on the literatures. The 32 ICRs were downloaded from the published work^55^.

### Partial correlation analysis of nucleosome occupancy and promoter NDR establishment

To quantify the relationship of newly established nucleosomes at each PN stage with other genomic features such as GC content, DNA methylation level and chromatin openness (defined using ATAC-seq peaks), and to correct the potential effect caused by nucleosome occupancy in previous stages, we performed partial correlation analysis on nucleosome occupancy for each stage. Briefly, we performed linear regression of the newly gained nucleosome occupancy at the current stage with the nucleosome occupancy at the previous stage using the lm function implemented in R. We also similarly performed linear regression of GC content or other features with the nucleosome occupancy at the previous stage. Then, we calculated the Pearson correlation coefficients of the residuals from the two regression models as the partial correlation between nucleosome occupancy and the input feature, such as GC content. The relationship of newly established promoter NDRs with other genomic features was similarly determined.

### K-means clustering of genes based on promoter NDR scores and analysis of variance (ANOVA)

To investigate the features correlated with promoter NDR scores, we first clustered the genes based on promoter NDR scores at 10 male PN stages using k-means clustering, setting k=7. Heatmap was generated using pheatmap function in R. We then performed ANNOVA analysis on 7 NDR clusters with histone modifications defined at sperm, zygote and 2-cell stages respectively as well as the GC content using aov functions in R, and the F-value of each histone modification was used to evaluate its association with NDR scores.

### Enrichment of ZGA gene promoters and GC content on TF binding sites

We calculated the enrichment of minor and major ZGA gene promoters on binding regions of different TFs as the odds ratio between the observed and expected counts. The observed count is calculated as the number of ZGA gene promoters containing the specific TF motif divided by the total number of promoters containing this motif. The expected count is calculated as the number of ZGA genes divided by the total number of genes. GC content on binding sites of a designated TF was calculated as the averaged GC content of top 10000 sites with highest motif scores across the genome.

### Differential expression analysis

To identify differentially expressed genes (DEGs), we calculated the read counts of each RNA-seq sample using HTSeq (v0.6.0)^56^. The counts in different replicates were fed into edgeR to perform differential expression analysis^57^. Genes with a p-value (Benjamini and Hochberg-adjusted) less than 0.05 and a fold change larger than 2 were defined as differentially expressed genes. Minor ZGA associated genes were defined as genes upregulated at the zygote stage compared to MII oocytes (1640 genes), and major ZGA associated genes were defined as genes upregulated at the 2-cell stage compared to zygotes (1012 genes) using our previously published RNA-seq data^58, 59^.

### Statistics and reproducibility

Error bars in the graphical data represent the standard deviation (SD). For all the presented boxplots, the center represents the median value, and the lower and upper lines represent the 5% and 95% quantiles, respectively. Significant difference between different groups was determined using the one-sided Wilcoxon rank-sum test adjusted by the Benjamini and Hochberg method^60^, and p < 0.05 was considered to be statistically significant. MNase-seq and RNA-seq experiments were performed two to five times for each group, and the precise numbers of replicates and the data qualities were summarized in supplementary Table S1. The information for MNase-seq sample normalization was provided in supplementary Table S2.

### Data availability

All the MNase-seq and RNA-seq data generated in this study were summarized in supplementary Table S1 and have been deposited to the GEO database under the accession number GSE140877. Sperm histone modification, ATAC-seq and DNA methylation data were downloaded from the GEO database (GSE79229)^23^. Zygote histone modification dataset were downloaded from GSE143523^61^. Bulk MNase-seq data of ESCs were downloaded from GSE51766^17^. TH2A, TH2B and input ChIP-seq data were downloaded from SRX398496^24^. RNA-seq data of mouse early embryos were downloaded from our previous publication (GSE97778)^59^. All the other data supporting the findings of this study are available from the corresponding author upon reasonable request.

## Acknowledgements

We thank Dr.K,Yamagata for providing the H2B-RFP plasmid. This work was supported by the National Key R&D Program of China (2016YFA0100400, 2018YFA0108900 and 2017YFA0102600), National Natural Science Foundation of China (31922022, 31721003, 31801059, 31771646, 32070802, 31970642, 32030022, 32170660), the Science and Technology Commission of Shanghai Municipality (19JC1415300, 17QA1404200), the Shanghai Municipal Medical and Health Discipline Construction Projects (2017ZZ02015), the Major Program of Development Fund for Shanghai Zhangjiang National Innovtaion Demonstration Zone (ZJ2018-ZD-004) and the Fundamental Research Funds for the Central Universities (1515219049). This work was also supported by the National Postdoctoral Program for Innovative Talents (BX20170174). Shanghai Rising Star Program [21QA1408200]. Natural Science Foundation of Shanghai [21ZR1467600]. The authors thank the Bioinformatics Supercomputer Center of Tongji University for offering computing resource.

## Author Contributions

Y.G., Y.Z. and S.G. conceived the project. Y.G. and S.G. designed the experiments. C.W. developed the NEPTUNE method and performed bioinformatic analyses. C.C., X.L. and C.L. performed the experiments. L.Y., X.K., Y.Z., and H.W assisted with the sample preparation. C.W., Y.G., C.C., X.L., Y.Z. and S.G. wrote the manuscript.

---

**Fig. S1 Development of ULI-MNase-seq with mESC samples. a** Schematic showing the procedures of ULI-MNase-seq. **b** Density plot showing the length distribution of mapped reads in mESC MNase-seq libraries started from different amounts of input. **c** Heatmap showing Pearson’s correlation coefficients between MNase-seq replicates of mESC samples from different amounts of input, which were calculated based on the genome-wide nucleosome signal. **d** Density plot showing the distribution of the relative nucleosome occupancy of 1-kb bins in mESC samples from different amounts of input. The dashed line represents the cut-off for nucleosome-occupied regions (occupancy O > 0.3). **e** Bar plots showing the fraction of nucleosome-occupied 1-kb bins in mESC samples from different amounts of input. Error bars represent ±1.96*SD. **f, g and h** Nucleosome profiles around TSSs of Refseq genes (f), enhancers (g; defined using ESC ATAC-seq peaks) or CTCF motifs (h) in replicates of mESC samples from different amounts of input.

**Fig. S2 NEPTUNE for integrated analysis of ULI-MNase-seq data. a** Schematic diagram showing the preprocessing steps for the NEPTUNE pipeline. After mapping raw nucleosome datasets to the genome and filtering low-quality reads, NEPTUNE performs tag extension and piling up, and then normalizes the nucleosome signal according to sequence depth. NEPTUNE applies signal correlation analysis on replicates, read length analysis, genome coverage analysis and nucleosome profiling around specific regions as quality controls of ULI-MNase-seq data. **b** NEPTUNE estimates the nucleosome occupancy by taking the genome coverage of nucleosome reads and background noise into account. NEPTUNE also calculates nucleosome depletion scores (NDR scores) and phasing scores (POS scores) for summarizing the nucleosome positioning pattern. **c** NEPTUNE generates nucleosome profiles around TSSs of all genes or specific gene sets and calculates corresponding NDR scores as well as the genome-wide nucleosome coverage to evaluate the influence of different perturbations on nucleosome positioning. **d** NEPTUNE analyzes the dynamic change of NDR pattern on motif regions for individual regulators, and also classifies multiple regulators based on the NDR dynamics.

**Fig. S3 Quality controls of ULI-MNase-seq in mouse pronuclei. a** Illustration of the parental pronuclei at 3 hpf, 6 hpf and 12 hpf with Hoechst staining. **b** Boxplots showing the Pearson’s correlation coefficients between MNase-seq replicates of each PN sample, which were calculated based on the nucleosome signal on promoter regions. **c** Scatter plots showing genome-wide nucleosome correlations between replicates of 1.5-hpf PN samples. **d** Density plots showing the length distribution of mapped reads in MNase-seq libraries at each PN stage. **e** Bar plots showing the fraction of short (5-50 bp) and long (120-180 bp) mapped reads in MNase-seq libraries at each PN stage. **f** Density plots showing the length distribution of mapped MNase-seq reads in sperm MNase-seq libraries under different conditions of MNase digestion. Color keys represent the amount of MNase used in each reaction, and subtitles indicate the duration of MNase digestion. **g** Bar plots showing the enrichment of short (5-50 bp) and long (120-180 bp) reads defined in sperm, 0.5-hpf or 1-hpf male PN on different genomic elements. **h** PCA analyses of ULI-MNase-seq samples at each PN stage. M, male PN. F, female PN. RSM, male PN of RS-fertilized embryos. RSF, female PN of RS-fertilized embryos. h, hpf.

**Fig. S4 Features of nucleosome occupancy in mouse pronuclei. a** Bar plots showing the enrichment of newly established nucleosome regions defined at each PN stage on different genomic elements. **b** UCSC genome browser view of a sperm-retained nucleosome locus (left) and a locus with nucleosomes established in 1-hpf male PN (right). **c** GO analysis of genes with nucleosomes retained in sperm or newly established in 1-hpf/6-hpf male PN. **d** Nucleosome profiles around TH2A/TH2B peaks in X chromosomes or other chromosomes at each PN stage. **e** Bar plots showing the Pearson’s correlation coefficients between nucleosome signals and TH2A/TH2B signals at each PN stage. h, hpf. Chr, chromosome.

**Fig. S5 Features of nucleosome positioning in mouse pronuclei. a** Graph showing the auto-correlation of nucleosome signals downstream of TSSs at each PN stage. Y-axis represents the Pearson’s correlation coefficient between the nucleosome signal of a designate site and the +10 bp site. Phasing periodicity (illustrated for 4-hpf male PN) represents the distance of the first peak summit from the TSS, and peaks within TSS +100 bp were ignored. h, hpf. **b** Nucleosome profiles around paternal and maternal imprinting control regions (ICRs) at each PN stage. **c** Boxplots showing NDR scores on promoters of all Refseq genes, maternally imprinted genes or paternally imprinted genes at each PN stage. **d and e** Nucleosome profiles around late 2-cell-defined (d) or ICM-defined (e) enhancers (defined using ATAC-seq peaks) at each PN stage. **f and g** Boxplots showing NDR scores (f) or POS scores (g) on promoters of all Refseq genes and ZGA genes at each PN stage. Significant difference is calculated between the designated gene set with all genes (*** p < 0.001; ** p < 0.01; * p < 0.05), and it is not labeled if p > 0.05.

**Fig. S6 Determinants of nucleosome occupancy and positioning in mouse pronuclei. a** Boxplots showing the GC content of newly established nucleosome regions at each PN stage after ROSI-mediated fertilization. Dashed lines represent the average GC content in genome. **b and c** Bar plots showing the partial correlation between nucleosome occupancy and GC content (b), chromatin accessibility (c left; defined using sperm ATAC-seq peaks), or DNA methylation (c right; defined using sperm whole-genome bisulfite sequencing data) at each PN stage. **d and e** Bar plots showing the partial correlation between promoter NDR scores and GC content (d), chromatin accessibility (e left; defined same with c left) or DNA methylation (e right; defined as c right) at each PN stage. **f** Heatmap showing the k-means clustering (k=7) of coding genes based on promoter NDR scores at each male PN stage. **g** Boxplots showing the level of indicated histone modifications on promoters of genes in different promoter NDR clusters (defined in f). Y-axis represents the normalized ChIP-seq signals. **h, i and j** Confocal microscopy images of 12-hpf treated embryos after EU (h and j) or EdU (i) staining. **k, l and m** Density plots showing the length distribution of mapped reads in MNase-seq libraries of 6-hpf or 12-hpf parental PN from groups under different treatment.

**Fig. S7 Histone acetylation influences NDR establishment in mouse male pronuclei. a and b** Scatter plots showing genome-wide nucleosome correlations between 6-hpf PN samples under different treatment. The first (a) and the second (b) batch of data were compared separately**. c and d** Boxplots showing the promoter NDR scores in 6-hpf (c) or 12-hpf (d) parental PN from groups under different treatment (*** p < 0.001; N.S. p > 0.05). **e, f and g** Nucleosome profiles around TSSs of ZGA genes in 6-hpf (e and g) or 12-hpf (f) parental PN from groups under different treatment.

**Fig. S8 Screening for potential ZGA-associated transcription factors. a and b** Nucleosome profiles around ARNTL (a) or POU5F1 (b) motifs at each PN stage. NDR scores on motif regions calculated based on the averaged nucleosome profile of each stage are labeled. **c** Boxplot showing the GC content of motifs of TFs in different clusters (defined in Fig. 4b; *** p < 0.001). **d** Bar plots showing the enrichment of ZGA gene promoters on motifs of individual cluster-3 TFs (defined in Fig. 4b). **e** Bar plots showing the expression level of potential ZGA-associated TFs during mouse preimplantation development. Error bars represent ±1.96*SD.

**Fig. S9 Failure of promoter NDR establishment in male PN after *Mlx* or *Rfx1* silencing. a and b** Nucleosome profiles around MLX (a) or RFX1 (b) motifs at each PN stage. NDR scores of individual stages are labeled. **c** Density plots showing the length distribution of mapped reads in MNase-seq libraries of 8-hpf parental PN from KD groups. **d and e** Nucleosome profiles around TSSs of different classes of genes in 8-hpf maternal PN from KD groups. Genes were classified according to whether motifs of MLX (d) or RFX1 (e) are present in the promoters. **f** UCSC genome browser view of an RFX1 motif-containing site with a decrease in +1 nucleosomes after *Rfx1* knockdown in both male and female PN. **g and h** Nucleosome profiles around TSSs of different classes of genes in 8-hpf parental PN from KD groups. Genes were classified according to whether they are minor (g) or major (h) ZGA genes.

**Fig. S10 *Mlx* or *Rfx1* silencing blocks the ZGA process in mouse embryos. a** Bar plots showing the expression level of siRNA-targeted genes in KD embryos (** p < 0.01; * p < 0.05). Error bars represent ±1.96*SD. **b** PCA analyses of RNA-seq replicates from KD groups. **c** Scatter plots showing the averaged expression level of genes (x-axis) and the fold change of genes upon KD (y-axis). Differentially expressed genes are labeled in red. **d** Bar plots showing counts of differentially expressed genes in KD embryos. **e** GO analysis of downregulated genes in *Mlx* or *Rfx1* KD 2-cell embryos. **f** Heatmap showing the significance of overlaps (calculated as p-values of the hypergeometric test) between downregulated genes in KD embryos and indicated gene sets. **g** Bar plot showing the overall 2-cell rates of KD groups. n=3 biological replicates with approximately 30 embryos each. **h** Sequence logo representing the deduced consensus motif of MLX, RFX1 and NFYA. **i** Venn diagram showing the overlap between genes with RFX1 motifs and genes with MLX motifs. **j** Venn diagram showing the overlap between major ZGA genes, genes with NFYA motifs and genes with RFX1 motifs.

## Supplementary Information

Fig. S1 to S10

Table S1: Summarize of mapped reads

Table S2: Normalization of ULI-MNase-seq data

## References

1. Clift, D. & Schuh, M. Restarting life: fertilization and the transition from meiosis to mitosis. Nat Rev Mol Cell Biol 14, 549–562, doi:10.1038/nrm3643 (2013).

2. Okada, Y. & Yamaguchi, K. Epigenetic modifications and reprogramming in paternal pronucleus: sperm, preimplantation embryo, and beyond. Cell Mol Life Sci 74, 1957–1967, doi:10.1007/s00018-016-2447-z (2017).

3. Lu, F. et al. Establishing Chromatin Regulatory Landscape during Mouse Preimplantation Development. Cell 165, 1375–1388, doi:10.1016/j.cell.2016.05.050 (2016).

4. Guo, F. et al. Single-cell multi-omics sequencing of mouse early embryos and embryonic stem cells. Cell Res 27, 967–988, doi:10.1038/cr.2017.82 (2017).

5. Iurlaro, M., von Meyenn, F. & Reik, W. DNA methylation homeostasis in human and mouse development. Curr Opin Genet Dev 43, 101–109, doi:10.1016/j.gde.2017.02.003 (2017).

6. Li, L. et al. Single-cell multi-omics sequencing of human early embryos. Nat Cell Biol 20, 847–858, doi:10.1038/s41556-018-0123-2 (2018).

7. Eckersley-Maslin, M. A., Alda-Catalinas, C. & Reik, W. Dynamics of the epigenetic landscape during the maternal-to-zygotic transition. Nat Rev Mol Cell Biol 19, 436–450, doi:10.1038/s41580-018-0008-z (2018).

8. Liu, B. et al. The landscape of RNA Pol II binding reveals a stepwise transition during ZGA. Nature 587, 139–144, doi:10.1038/s41586-020-2847-y (2020).

9. Struhl, K. & Segal, E. Determinants of nucleosome positioning. Nat Struct Mol Biol 20, 267–273, doi:10.1038/nsmb.2506 (2013).

10. Chereji, R. V. & Clark, D. J. Major Determinants of Nucleosome Positioning. Biophys J 114, 2279–2289, doi:10.1016/j.bpj.2018.03.015 (2018).

11. Hughes, A. L. & Rando, O. J. Mechanisms underlying nucleosome positioning in vivo. Annu Rev Biophys 43, 41–63, doi:10.1146/annurev-biophys-051013-023114 (2014).

12. Maehara, K. & Ohkawa, Y. Exploration of nucleosome positioning patterns in transcription factor function. Sci Rep 6, 19620, doi:10.1038/srep19620 (2016).

13. Rhee, H. S. & Pugh, B. F. Genome-wide structure and organization of eukaryotic pre-initiation complexes. Nature 483, 295–301, doi:10.1038/nature10799 (2012).

14. Bai, L., Ondracka, A. & Cross, F. R. Multiple sequence-specific factors generate the nucleosome-depleted region on CLN2 promoter. Mol Cell 42, 465–476, doi:10.1016/j.molcel.2011.03.028 (2011).

15. Carone, B. R. et al. High-resolution mapping of chromatin packaging in mouse embryonic stem cells and sperm. Dev Cell 30, 11–22, doi:10.1016/j.devcel.2014.05.024 (2014).

16. Luger, K., Mader, A. W., Richmond, R. K., Sargent, D. F. & Richmond, T. J. Crystal structure of the nucleosome core particle at 2.8 A resolution. Nature 389, 251–260, doi:10.1038/38444 (1997).

17. Zhang, Y. et al. Canonical nucleosome organization at promoters forms during genome activation. Genome Res 24, 260–266, doi:10.1101/gr.157750.113 (2014).

18. Adenot, P. G., Mercier, Y., Renard, J. P. & Thompson, E. M. Differential H4 acetylation of paternal and maternal chromatin precedes DNA replication and differential transcriptional activity in pronuclei of 1-cell mouse embryos. Development 124, 4615–4625 (1997).

19. Kurotaki, Y. K. et al. Impaired active DNA demethylation in zygotes generated by round spermatid injection. Hum Reprod 30, 1178–1187, doi:10.1093/humrep/dev039 (2015).

20. Govin, J. et al. Pericentric heterochromatin reprogramming by new histone variants during mouse spermiogenesis. J Cell Biol 176, 283–294, doi:10.1083/jcb.200604141 (2007).

21. Zhou, L., Baibakov, B., Canagarajah, B., Xiong, B. & Dean, J. Genetic mosaics and time-lapse imaging identify functions of histone H3.3 residues in mouse oocytes and embryos. Development 144, 519–528, doi:10.1242/dev.141390 (2017).

22. Samans, B. et al. Uniformity of nucleosome preservation pattern in Mammalian sperm and its connection to repetitive DNA elements. Dev Cell 30, 23–35, doi:10.1016/j.devcel.2014.05.023 (2014).

23. Jung, Y. H. et al. Chromatin States in Mouse Sperm Correlate with Embryonic and Adult Regulatory Landscapes. Cell Rep 18, 1366–1382, doi:10.1016/j.celrep.2017.01.034 (2017).

24. Shinagawa, T. et al. Histone variants enriched in oocytes enhance reprogramming to induced pluripotent stem cells. Cell Stem Cell 14, 217–227, doi:10.1016/j.stem.2013.12.015 (2014).

25. Wu, J. et al. The landscape of accessible chromatin in mammalian preimplantation embryos. Nature 534, 652–657, doi:10.1038/nature18606 (2016).

26. Aoki, F., Worrad, D. M. & Schultz, R. M. Regulation of transcriptional activity during the first and second cell cycles in the preimplantation mouse embryo. Dev Biol 181, 296–307, doi:10.1006/dbio.1996.8466 (1997).

27. Huff, J. T. & Zilberman, D. Dnmt1-independent CG methylation contributes to nucleosome positioning in diverse eukaryotes. Cell 156, 1286–1297, doi:10.1016/j.cell.2014.01.029 (2014).

28. Zhang, Z. et al. A packing mechanism for nucleosome organization reconstituted across a eukaryotic genome. Science 332, 977–980, doi:10.1126/science.1200508 (2011).

29. Zhang, Y. et al. Intrinsic histone-DNA interactions are not the major determinant of nucleosome positions in vivo. Nat Struct Mol Biol 16, 847–852, doi:10.1038/nsmb.1636 (2009).

30. Filippakopoulos, P. et al. Selective inhibition of BET bromodomains. Nature 468, 1067–1073, doi:10.1038/nature09504 (2010).

31. Jung, M. et al. Affinity map of bromodomain protein 4 (BRD4) interactions with the histone H4 tail and the small molecule inhibitor JQ1. J Biol Chem 289, 9304–9319, doi:10.1074/jbc.M113.523019 (2014).

32. Workman, J. L. & Kingston, R. E. Nucleosome core displacement in vitro via a metastable transcription factor-nucleosome complex. Science 258, 1780–1784, doi:10.1126/science.1465613 (1992).

33. Jiang, C. & Pugh, B. F. Nucleosome positioning and gene regulation: advances through genomics. Nat Rev Genet 10, 161–172, doi:10.1038/nrg2522 (2009).

34. Chen, X. et al. Key role for CTCF in establishing chromatin structure in human embryos. Nature, doi:10.1038/s41586-019-1812-0 (2019).

35. Mei, S. et al. Cistrome Data Browser: a data portal for ChIP-Seq and chromatin accessibility data in human and mouse. Nucleic acids research 45, D658–D662, doi:10.1093/nar/gkw983 (2017).

36. Hunt, L. C. et al. The glucose-sensing transcription factor MLX promotes myogenesis via myokine signaling. Genes Dev 29, 2475–2489, doi:10.1101/gad.267419.115 (2015).

37. Safrany, G. & Perry, R. P. Transcription factor RFX1 helps control the promoter of the mouse ribosomal protein-encoding gene rpL30 by binding to its alpha element. Gene 132, 279–283, doi:10.1016/0378-1119(93)90208-k (1993).

38. Ma, K., Zheng, S. & Zuo, Z. The transcription factor regulatory factor X1 increases the expression of neuronal glutamate transporter type 3. J Biol Chem 281, 21250–21255, doi:10.1074/jbc.M600521200 (2006).

39. Wang, B. et al. RFX1 maintains testis cord integrity by regulating the expression of Itga6 in male mouse embryos. Mol Reprod Dev 83, 606–614, doi:10.1002/mrd.22660 (2016).

40. Mizukawa, T. et al. RFX1-mediated CCN3 induction that may support chondrocyte survival under starved conditions. J Cell Physiol 236, 6884–6896, doi:10.1002/jcp.30348 (2021).

41. Feng, C., Xu, W. & Zuo, Z. Knockout of the regulatory factor X1 gene leads to early embryonic lethality. Biochem Biophys Res Commun 386, 715–717, doi:10.1016/j.bbrc.2009.06.111 (2009).

42. Hajkova, P. Epigenetic reprogramming--taking a lesson from the embryo. Curr Opin Cell Biol 22, 342–350, doi:10.1016/j.ceb.2010.04.011 (2010).

43. Vastenhouw, N. L., Cao, W. X. & Lipshitz, H. D. The maternal-to-zygotic transition revisited. Development 146, doi:10.1242/dev.161471 (2019).

44. Soufi, A. et al. Pioneer transcription factors target partial DNA motifs on nucleosomes to initiate reprogramming. Cell 161, 555–568, doi:10.1016/j.cell.2015.03.017 (2015).

45. Picelli, S. et al. Full-length RNA-seq from single cells using Smart-seq2. Nat Protoc 9, 171–181, doi:10.1038/nprot.2014.006 (2014).

46. Li, H. & Durbin, R. Fast and accurate long-read alignment with Burrows-Wheeler transform. Bioinformatics 26, 589–595, doi:10.1093/bioinformatics/btp698 (2010).

47. Quinlan, A. R. & Hall, I. M. BEDTools: a flexible suite of utilities for comparing genomic features. Bioinformatics 26, 841–842, doi:10.1093/bioinformatics/btq033 (2010).

48. Shin, H., Liu, T., Manrai, A. K. & Liu, X. S. CEAS: cis-regulatory element annotation system. Bioinformatics 25, 2605–2606, doi:10.1093/bioinformatics/btp479 (2009).

49. Zhang, Y. et al. Model-based analysis of ChIP-Seq (MACS). Genome Biol 9, R137, doi:10.1186/gb-2008-9-9-r137 (2008).

50. Dobin, A. et al. STAR: ultrafast universal RNA-seq aligner. Bioinformatics 29, 15–21, doi:10.1093/bioinformatics/bts635 (2013).

51. Pertea, M. et al. StringTie enables improved reconstruction of a transcriptome from RNA-seq reads. Nature biotechnology 33, 290–295, doi:10.1038/nbt.3122 (2015).

52. Wang, B. et al. Integrative analysis of pooled CRISPR genetic screens using MAGeCKFlute. Nat Protoc 14, 756–780, doi:10.1038/s41596-018-0113-7 (2019).

53. Babak, T. et al. Genetic conflict reflected in tissue-specific maps of genomic imprinting in human and mouse. Nat Genet 47, 544–549, doi:10.1038/ng.3274 (2015).

54. Wei, Y. et al. MetaImprint: an information repository of mammalian imprinted genes. Development 141, 2516–2523, doi:10.1242/dev.105320 (2014).

55. Xie, W. et al. Base-resolution analyses of sequence and parent-of-origin dependent DNA methylation in the mouse genome. Cell 148, 816–831, doi:10.1016/j.cell.2011.12.035 (2012).

56. Anders, S., Pyl, P. T. & Huber, W. HTSeq--a Python framework to work with high-throughput sequencing data. Bioinformatics 31, 166–169, doi:10.1093/bioinformatics/btu638 (2015).

57. Robinson, M. D., McCarthy, D. J. & Smyth, G. K. edgeR: a Bioconductor package for differential expression analysis of digital gene expression data. Bioinformatics 26, 139–140, doi:10.1093/bioinformatics/btp616 (2010).

58. Liu, W. et al. Identification of key factors conquering developmental arrest of somatic cell cloned embryos by combining embryo biopsy and single-cell sequencing. Cell Discov 2, 16010, doi:10.1038/celldisc.2016.10 (2016).

59. Wang, C. et al. Reprogramming of H3K9me3-dependent heterochromatin during mammalian embryo development. Nat Cell Biol 20, 620–631, doi:10.1038/s41556-018-0093-4 (2018).

60. Jafari, M. & Ansari-Pour, N. Why, When and How to Adjust Your P Values? Cell J 20, 604–607, doi:10.22074/cellj.2019.5992 (2019).

61. Yang, G. et al. Dux-Mediated Corrections of Aberrant H3K9ac during 2-Cell Genome Activation Optimize Efficiency of Somatic Cell Nuclear Transfer. Cell Stem Cell 28, 150–163 e155, doi:10.1016/j.stem.2020.09.006 (2021).

