## Supplementary Figures for "Dynamic nucleosome organization after fertilization reveals regulatory factors for mouse zygotic genome activation"

### Figure S1

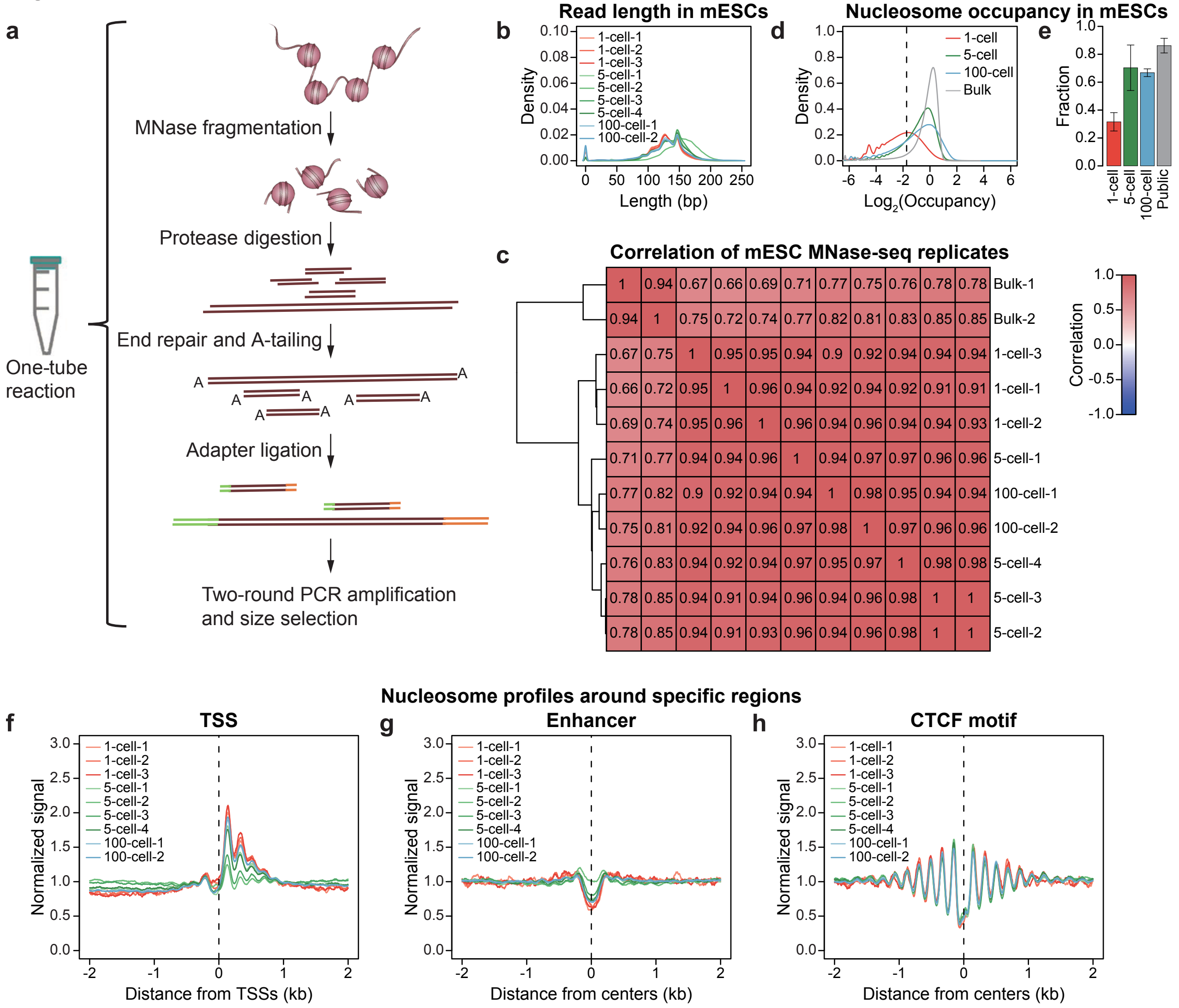

a
MNase fragmentation
Protease digestion
End repair and A-tailing
Adapter ligation
Two-round PCR amplification and size selection
One-tube reaction
b
Read length in mESCs
Density
Length (bp)
1-cell-1
1-cell-2
1-cell-3
5-cell-1
5-cell-2
5-cell-3
5-cell-4
100-cell-1
100-cell-2
d
Nucleosome occupancy in mESCs
Density
Log2(Occupancy)
1-cell
5-cell
100-cell
Bulk
e
Fraction
1-cell
5-cell
100-cell
Public
c
Correlation of mESC MNase-seq replicates
Correlation
Bulk-1
Bulk-2
1-cell-3
1-cell-1
1-cell-2
5-cell-1
100-cell-1
100-cell-2
5-cell-4
5-cell-3
5-cell-2
1 0.94 0.67 0.66 0.69 0.71 0.77 0.75 0.76 0.78 0.78
0.94 1 0.75 0.72 0.74 0.77 0.82 0.81 0.83 0.85 0.85
0.67 0.75 1 0.95 0.95 0.94 0.9 0.92 0.94 0.94 0.94
0.66 0.72 0.95 1 0.96 0.94 0.92 0.94 0.92 0.91 0.91
0.69 0.74 0.95 0.96 1 0.96 0.94 0.96 0.94 0.94 0.93
0.71 0.77 0.94 0.94 0.96 1 0.94 0.97 0.97 0.96 0.96
0.77 0.82 0.9 0.92 0.94 0.94 1 0.98 0.95 0.94 0.94
0.75 0.81 0.92 0.94 0.96 0.97 0.98 1 0.97 0.96 0.96
0.76 0.83 0.94 0.92 0.94 0.97 0.95 0.97 1 0.98 0.98
0.78 0.85 0.94 0.91 0.94 0.96 0.94 0.96 0.98 1 1
0.78 0.85 0.94 0.91 0.93 0.96 0.94 0.96 0.98 1 1
Nucleosome profiles around specific regions
f
TSS
Normalized signal
Distance from TSSs (kb)
g
Enhancer
Distance from centers (kb)
h
CTCF motif
Distance from centers (kb)

Figure S2

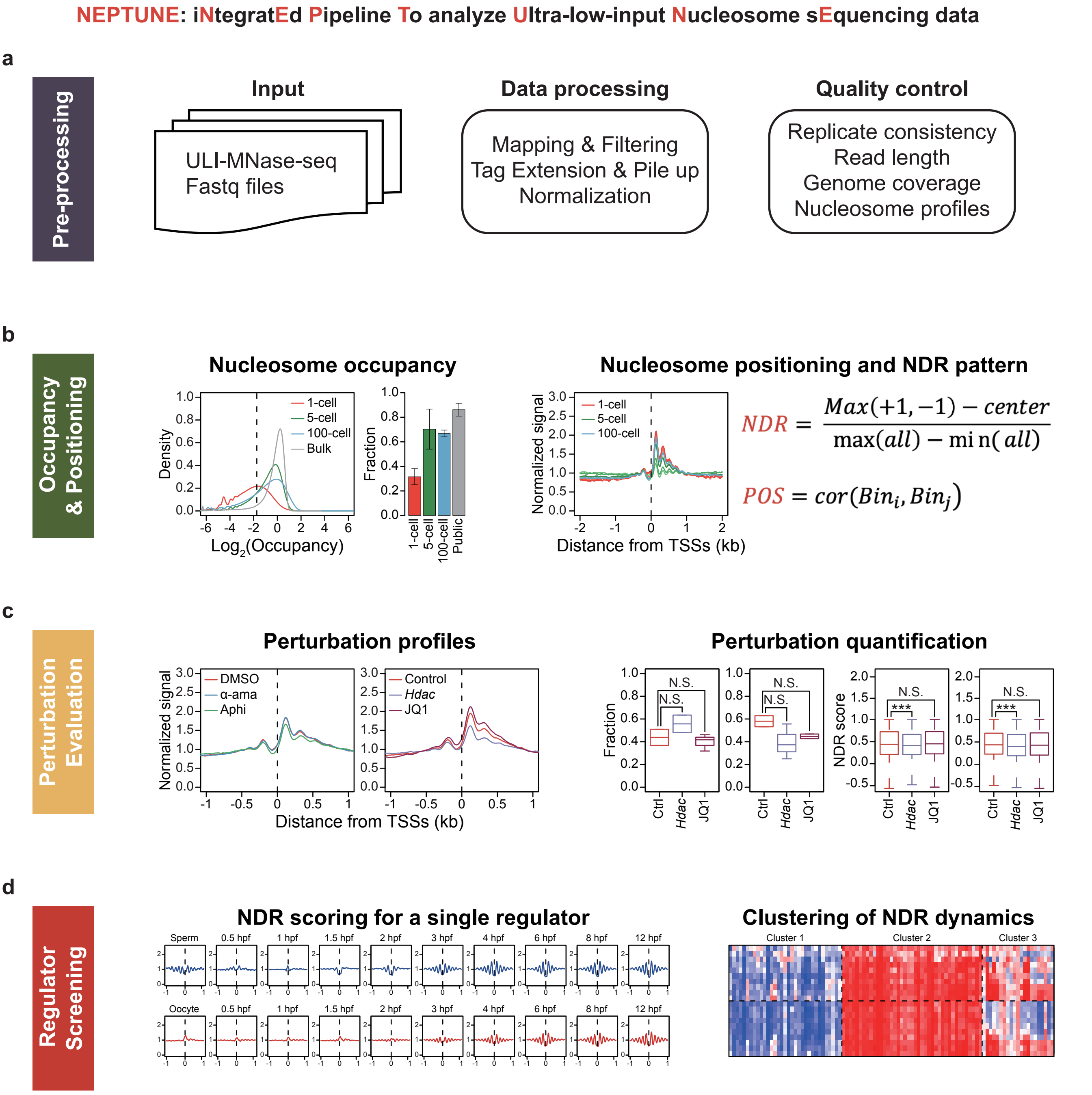
NEPTUNE: iNtegratEd Pipeline To analyze Ultra-low-input Nucleosome sEquencing data
a
Pre-processing
Input
ULI-MNase-seq
Fastq files
Data processing
Mapping & Filtering
Tag Extension & Pile up
Normalization
Quality control
Replicate consistency
Read length
Genome coverage
Nucleosome profiles
b
Occupancy & Positioning
Nucleosome occupancy
Density
Log2(Occupancy)
1-cell
5-cell
100-cell
Bulk
Fraction
Public
Nucleosome positioning and NDR pattern
Normalized signal
Distance from TSSs (kb)
NDR = (Max(+1, −1) − center) / (max(all) − min(all))
POS = cor(Bin_i, Bin_j)
c
Perturbation Evaluation
Perturbation profiles
DMSO
α-ama
Aphi
Control
Hdac
JQ1
Normalized signal
Distance from TSSs (kb)
Perturbation quantification
Fraction
NDR score
N.S.
***
Ctrl
Hdac
JQ1
d
Regulator Screening
NDR scoring for a single regulator
Sperm
Oocyte
0.5 hpf
1 hpf
1.5 hpf
2 hpf
3 hpf
4 hpf
6 hpf
8 hpf
12 hpf
Clustering of NDR dynamics
Cluster 1
Cluster 2
Cluster 3

Figure S3

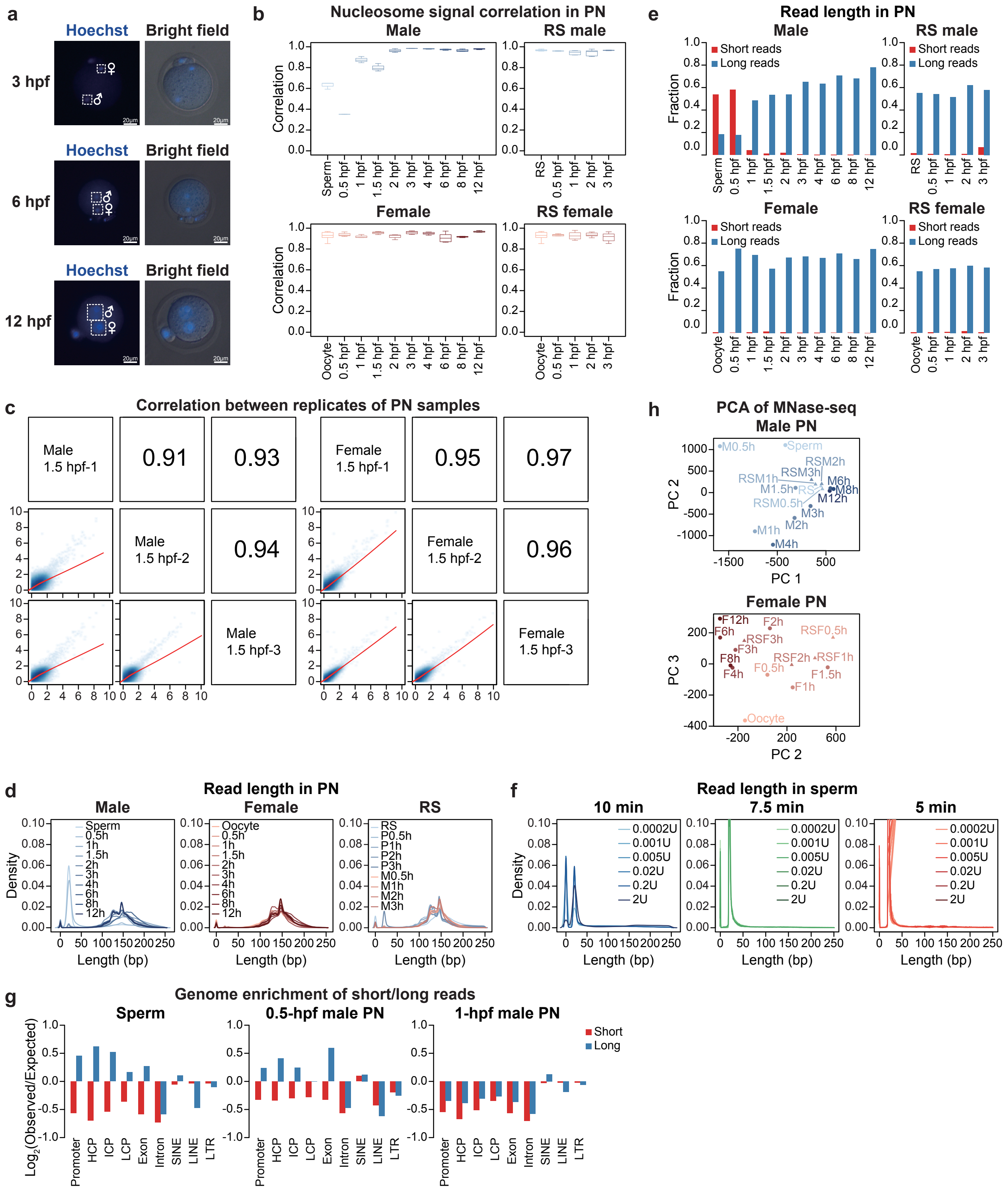
a
Hoechst
Bright field
3 hpf
6 hpf
12 hpf
b
Nucleosome signal correlation in PN
Male
RS male
Female
RS female
Correlation
c
Correlation between replicates of PN samples
Male 1.5 hpf-1
Male 1.5 hpf-2
Male 1.5 hpf-3
Female 1.5 hpf-1
Female 1.5 hpf-2
Female 1.5 hpf-3
0.91
0.93
0.94
0.95
0.97
0.96
d
Read length in PN
Male
Female
RS
Density
Length (bp)
e
Read length in PN
Male
RS male
Female
RS female
Short reads
Long reads
Fraction
f
Read length in sperm
10 min
7.5 min
5 min
Density
Length (bp)
g
Genome enrichment of short/long reads
Sperm
0.5-hpf male PN
1-hpf male PN
Short
Long
Log2(Observed/Expected)
Promoter
HCP
ICP
LCP
Exon
Intron
SINE
LINE
LTR
h
PCA of MNase-seq
Male PN
Female PN
PC 1
PC 2
PC 3

### Figure S4

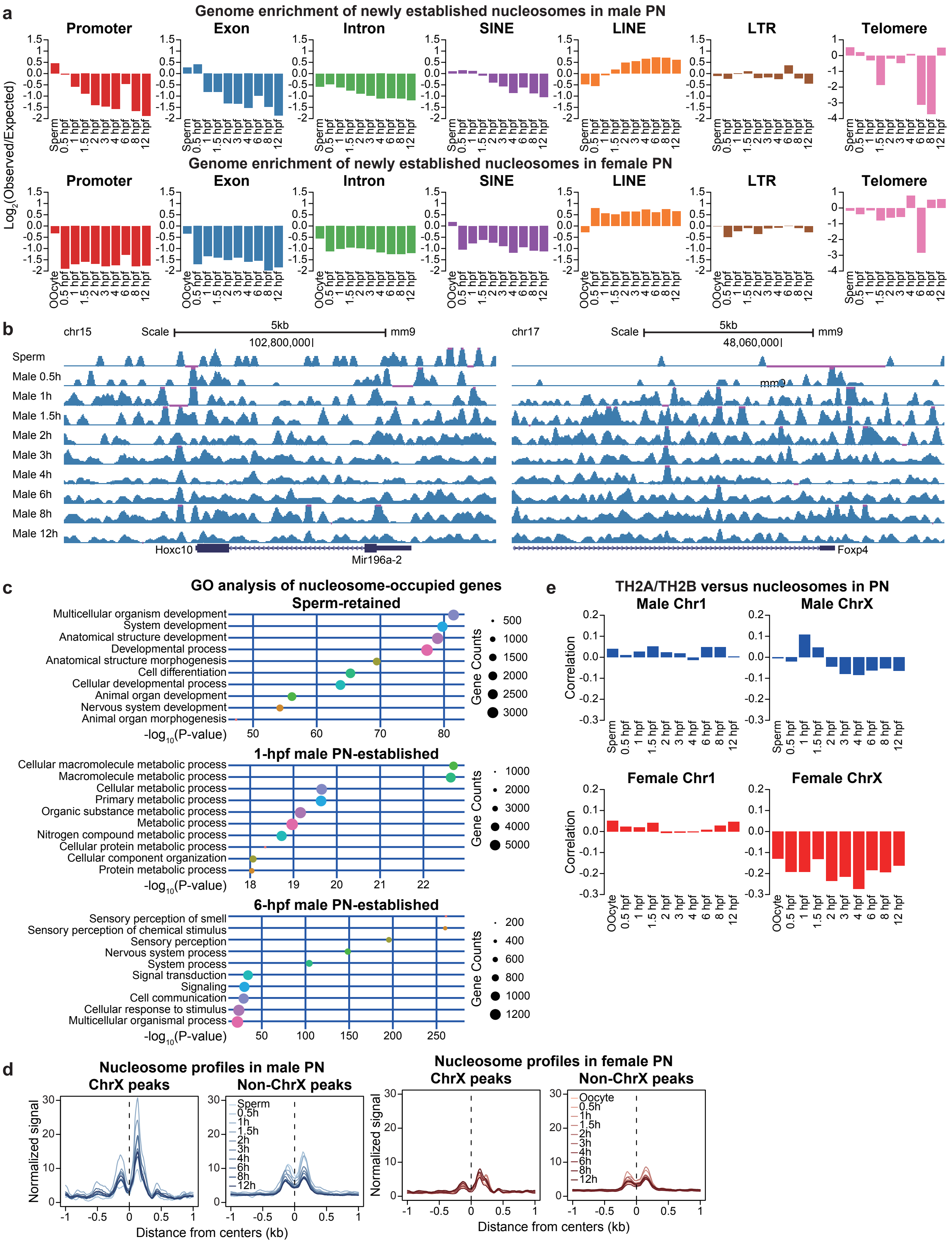

a
Genome enrichment of newly established nucleosomes in male PN
Promoter
Exon
Intron
SINE
LINE
LTR
Telomere
Log2(Observed/Expected)
Sperm
0.5 hpf
1 hpf
1.5 hpf
2 hpf
3 hpf
4 hpf
6 hpf
8 hpf
12 hpf
Genome enrichment of newly established nucleosomes in female PN
OOcyte
b
chr15
Scale
5kb
mm9
102,800,000|
chr17
48,060,000|
Sperm
Male 0.5h
Male 1h
Male 1.5h
Male 2h
Male 3h
Male 4h
Male 6h
Male 8h
Male 12h
Hoxc10
Mir196a-2
Foxp4
c
GO analysis of nucleosome-occupied genes
Sperm-retained
Multicellular organism development
System development
Anatomical structure development
Developmental process
Anatomical structure morphogenesis
Cell differentiation
Cellular developmental process
Animal organ development
Nervous system development
Animal organ morphogenesis
-log10(P-value)
Gene Counts
1-hpf male PN-established
Cellular macromolecule metabolic process
Macromolecule metabolic process
Cellular metabolic process
Primary metabolic process
Organic substance metabolic process
Metabolic process
Nitrogen compound metabolic process
Cellular protein metabolic process
Cellular component organization
Protein metabolic process
6-hpf male PN-established
Sensory perception of smell
Sensory perception of chemical stimulus
Sensory perception
Nervous system process
System process
Signal transduction
Signaling
Cell communication
Cellular response to stimulus
Multicellular organismal process
d
Nucleosome profiles in male PN
ChrX peaks
Non-ChrX peaks
Normalized signal
Distance from centers (kb)
Nucleosome profiles in female PN
Sperm
0.5h
1h
1.5h
2h
3h
4h
6h
8h
12h
Oocyte
e
TH2A/TH2B versus nucleosomes in PN
Male Chr1
Male ChrX
Female Chr1
Female ChrX
Correlation

Figure S5

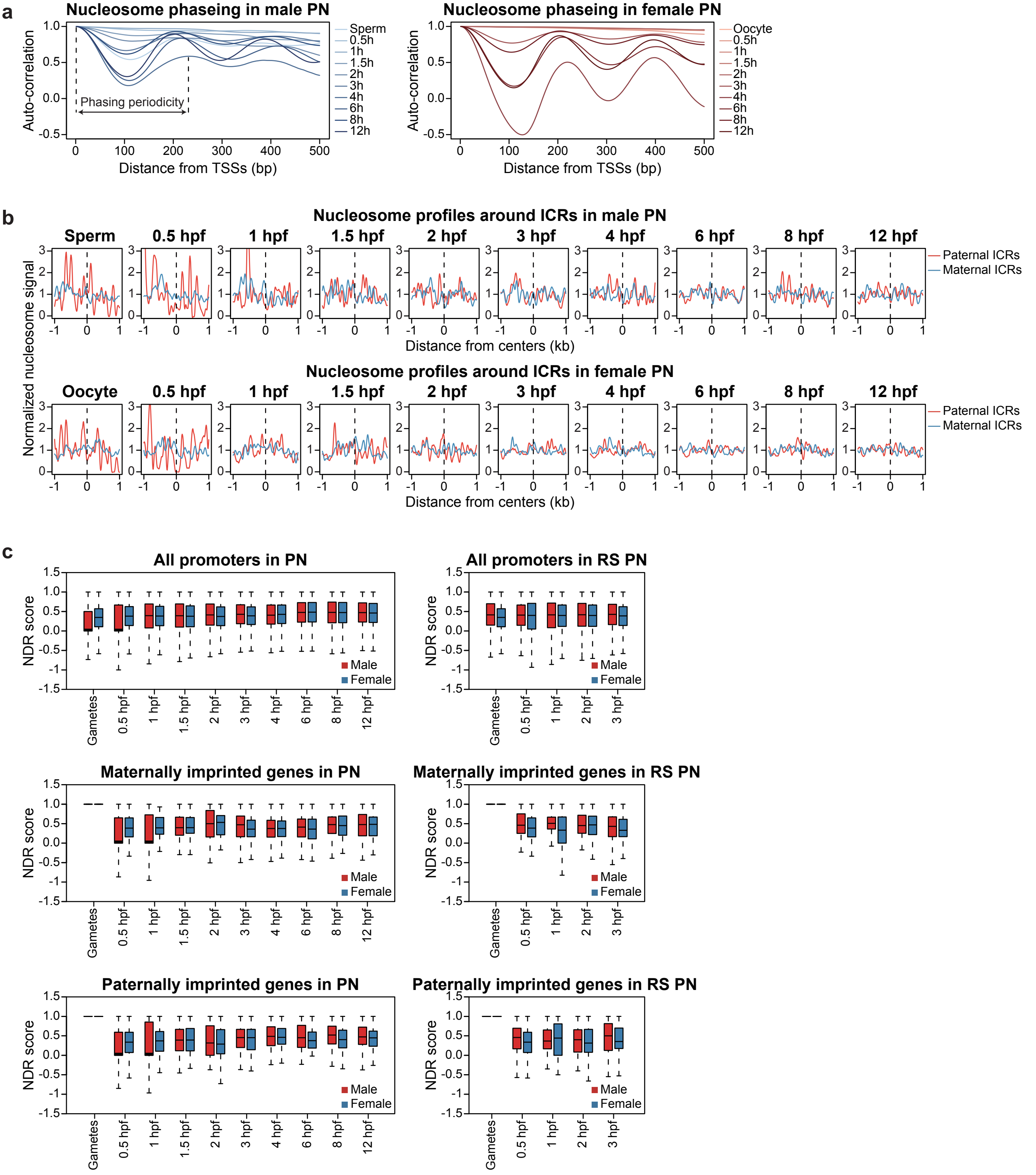


*(Continued on next page)*

*(Continued)*

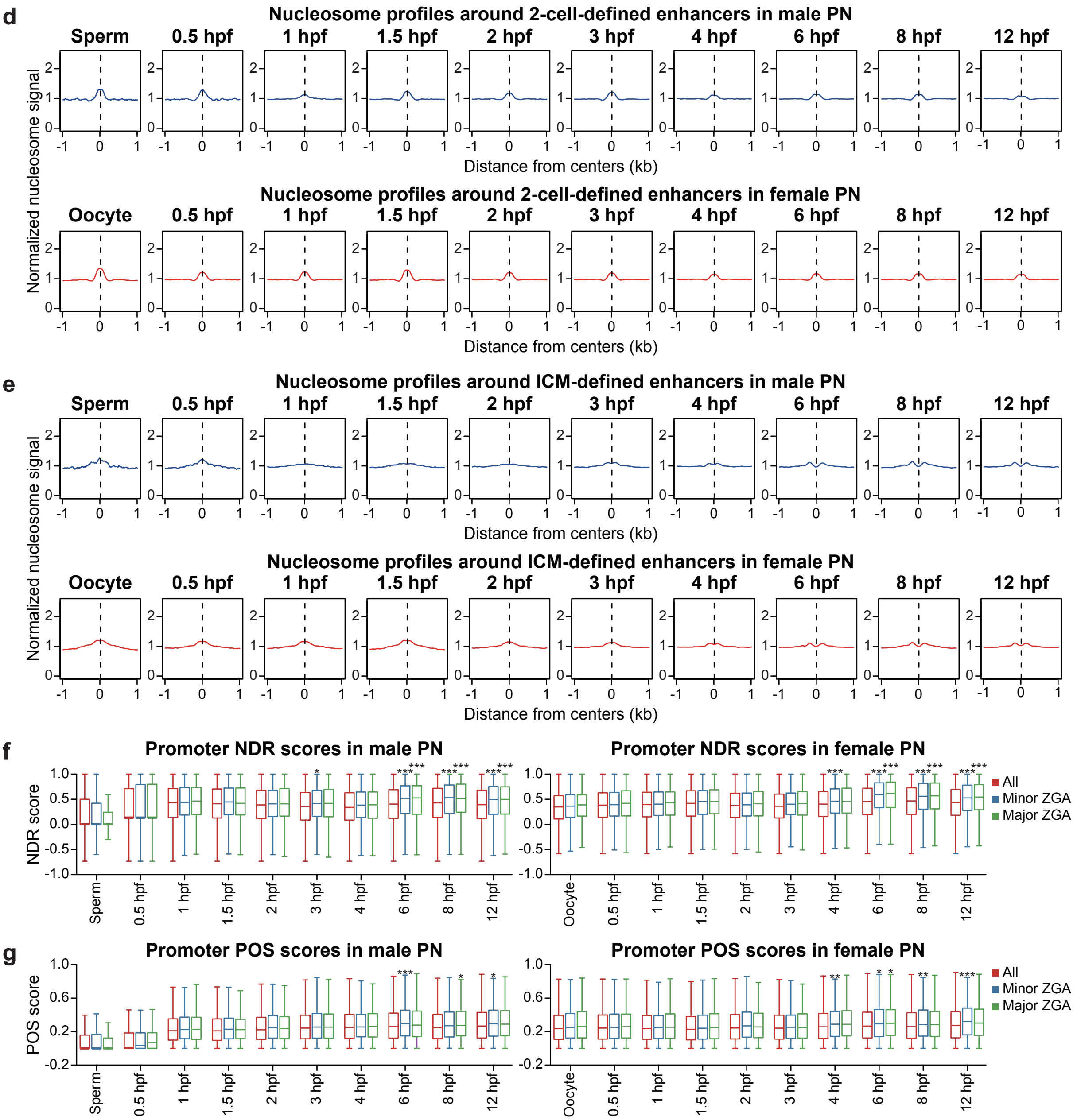
d
Nucleosome profiles around 2-cell-defined enhancers in male PN
Sperm
0.5 hpf
1 hpf
1.5 hpf
2 hpf
3 hpf
4 hpf
6 hpf
8 hpf
12 hpf
Normalized nucleosome signal
Distance from centers (kb)
Nucleosome profiles around 2-cell-defined enhancers in female PN
Oocyte
e
Nucleosome profiles around ICM-defined enhancers in male PN
Nucleosome profiles around ICM-defined enhancers in female PN
f
Promoter NDR scores in male PN
Promoter NDR scores in female PN
NDR score
All
Minor ZGA
Major ZGA
g
Promoter POS scores in male PN
Promoter POS scores in female PN
POS score

Figure S6

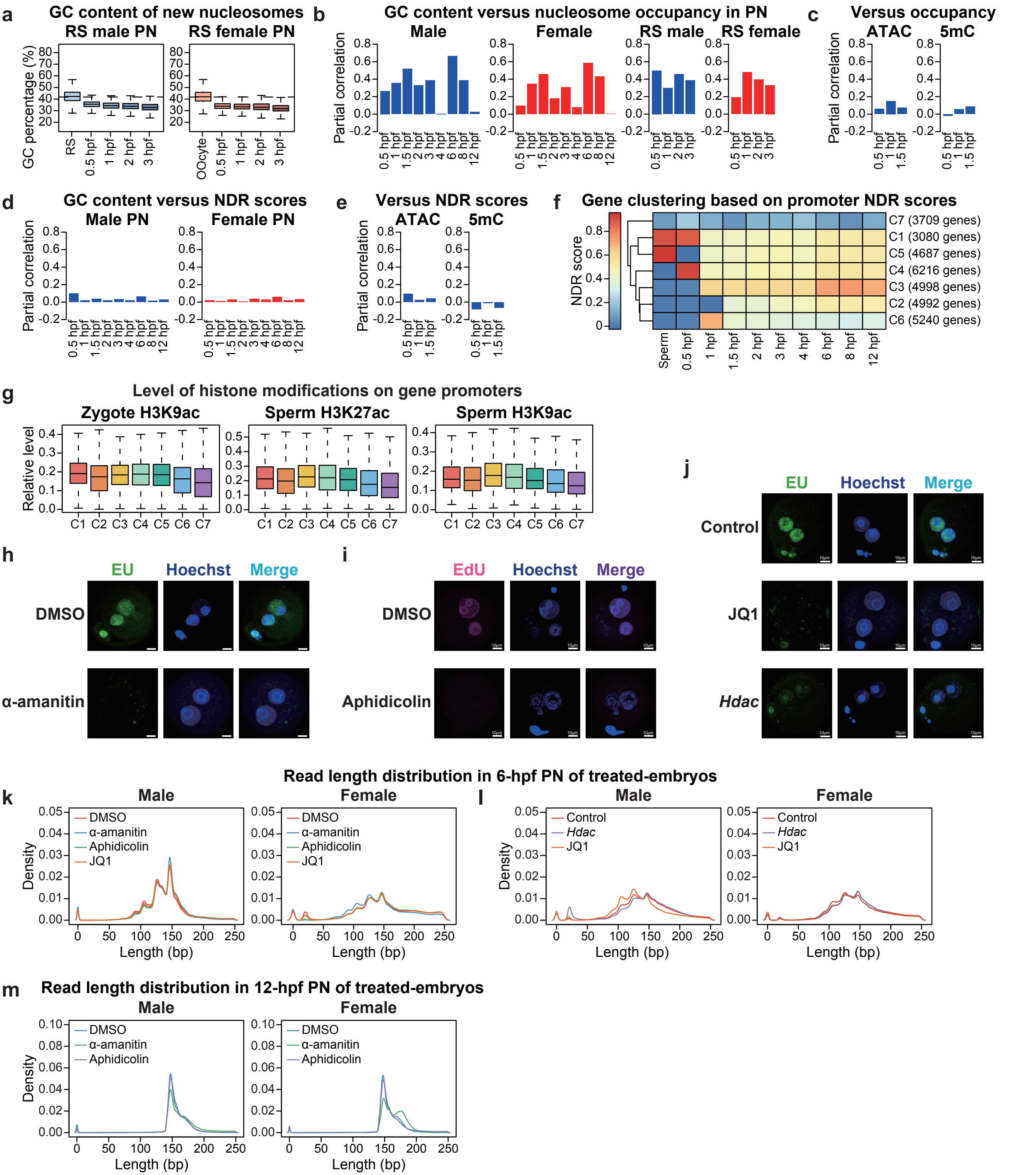
a
GC content of new nucleosomes
RS male PN
RS female PN
GC percentage (%)
RS
Oocyte
0.5 hpf
1 hpf
2 hpf
3 hpf
b
GC content versus nucleosome occupancy in PN
Male
Female
RS male
RS female
Partial correlation
c
Versus occupancy
ATAC
5mC
d
GC content versus NDR scores
Male PN
Female PN
e
Versus NDR scores
f
Gene clustering based on promoter NDR scores
NDR score
C7 (3709 genes)
C1 (3080 genes)
C5 (4687 genes)
C4 (6216 genes)
C3 (4998 genes)
C2 (4992 genes)
C6 (5240 genes)
Sperm
g
Level of histone modifications on gene promoters
Zygote H3K9ac
Sperm H3K27ac
Sperm H3K9ac
Relative level
h
EU
Hoechst
Merge
DMSO
α-amanitin
i
EdU
Aphidicolin
j
Control
JQ1
Hdac
Read length distribution in 6-hpf PN of treated-embryos
k
Male
Female
Density
Length (bp)
DMSO
α-amanitin
Aphidicolin
JQ1
l
Control
Hdac
m
Read length distribution in 12-hpf PN of treated-embryos

Figure S7

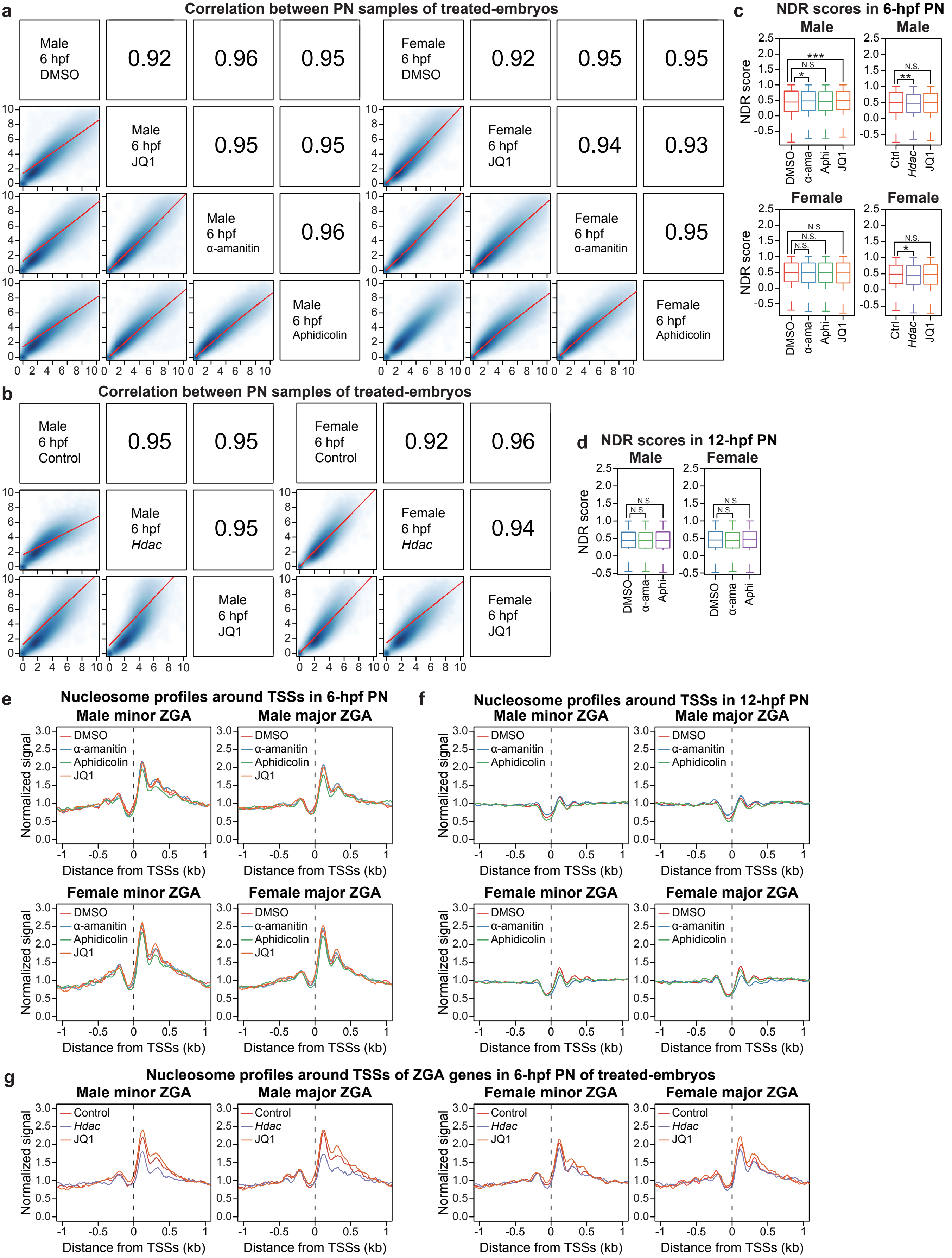
a
Correlation between PN samples of treated-embryos
Male 6 hpf DMSO
Male 6 hpf JQ1
Male 6 hpf α-amanitin
Male 6 hpf Aphidicolin
0.92
0.96
0.95
0.95
0.95
0.96
Female 6 hpf DMSO
Female 6 hpf JQ1
Female 6 hpf α-amanitin
Female 6 hpf Aphidicolin
0.92
0.95
0.95
0.94
0.93
0.95
b
Correlation between PN samples of treated-embryos
Male 6 hpf Control
Male 6 hpf Hdac
Male 6 hpf JQ1
0.95
0.95
0.95
Female 6 hpf Control
Female 6 hpf Hdac
Female 6 hpf JQ1
0.92
0.96
0.94
c
NDR scores in 6-hpf PN
Male
Female
NDR score
DMSO
α-ama
Aphi
JQ1
Ctrl
Hdac
N.S.
*
**
***
d
NDR scores in 12-hpf PN
Male
Female
NDR score
DMSO
α-ama
Aphi
N.S.
e
Nucleosome profiles around TSSs in 6-hpf PN
Male minor ZGA
Male major ZGA
Female minor ZGA
Female major ZGA
DMSO
α-amanitin
Aphidicolin
JQ1
Normalized signal
Distance from TSSs (kb)
f
Nucleosome profiles around TSSs in 12-hpf PN
Male minor ZGA
Male major ZGA
Female minor ZGA
Female major ZGA
DMSO
α-amanitin
Aphidicolin
Normalized signal
Distance from TSSs (kb)
g
Nucleosome profiles around TSSs of ZGA genes in 6-hpf PN of treated-embryos
Male minor ZGA
Male major ZGA
Female minor ZGA
Female major ZGA
Control
Hdac
JQ1
Normalized signal
Distance from TSSs (kb)

### Figure S8

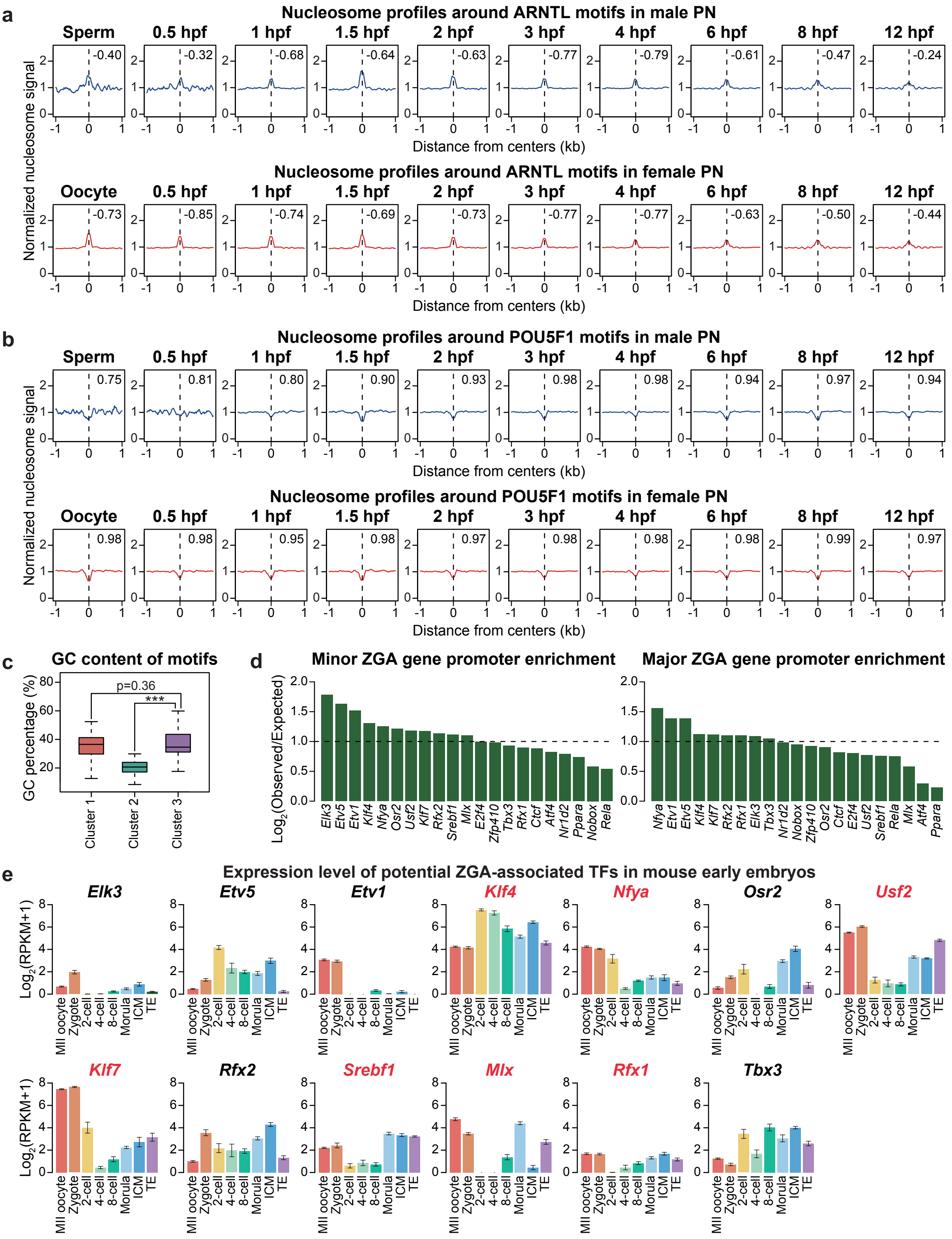
a
Nucleosome profiles around ARNTL motifs in male PN
Sperm 0.5 hpf 1 hpf 1.5 hpf 2 hpf 3 hpf 4 hpf 6 hpf 8 hpf 12 hpf
-0.40 -0.32 -0.68 -0.64 -0.63 -0.77 -0.79 -0.61 -0.47 -0.24
Normalized nucleosome signal
Distance from centers (kb)
Nucleosome profiles around ARNTL motifs in female PN
Oocyte 0.5 hpf 1 hpf 1.5 hpf 2 hpf 3 hpf 4 hpf 6 hpf 8 hpf 12 hpf
-0.73 -0.85 -0.74 -0.69 -0.73 -0.77 -0.77 -0.63 -0.50 -0.44
Distance from centers (kb)
b
Nucleosome profiles around POU5F1 motifs in male PN
Sperm 0.5 hpf 1 hpf 1.5 hpf 2 hpf 3 hpf 4 hpf 6 hpf 8 hpf 12 hpf
0.75 0.81 0.80 0.90 0.93 0.98 0.98 0.94 0.97 0.94
Normalized nucleosome signal
Distance from centers (kb)
Nucleosome profiles around POU5F1 motifs in female PN
Oocyte 0.5 hpf 1 hpf 1.5 hpf 2 hpf 3 hpf 4 hpf 6 hpf 8 hpf 12 hpf
0.98 0.98 0.95 0.98 0.97 0.98 0.98 0.98 0.99 0.97
Distance from centers (kb)
c
GC content of motifs
GC percentage (%)
p=0.36
***
Cluster 1 Cluster 2 Cluster 3
d
Minor ZGA gene promoter enrichment
Log2(Observed/Expected)
Elk3 Etv5 Etv1 Klf4 Nfya Osr2 Usf2 Klf7 Rfx2 Srebf1 Mlx E2f4 Zfp410 Tbx3 Rfx1 Ctcf Atf4 Nr1d2 Ppara Nobox Rela
Major ZGA gene promoter enrichment
Nfya Etv1 Etv5 Klf4 Klf7 Rfx2 Rfx1 Elk3 Tbx3 Nr1d2 Nobox Zfp410 Osr2 Ctcf E2f4 Usf2 Srebf1 Rela Mlx Atf4 Ppara
e
Expression level of potential ZGA-associated TFs in mouse early embryos
Log2(RPKM+1)
Elk3 Etv5 Etv1 Klf4 Nfya Osr2 Usf2
Klf7 Rfx2 Srebf1 Mlx Rfx1 Tbx3
MII oocyte Zygote 2-cell 4-cell 8-cell Morula ICM TE

Figure S9

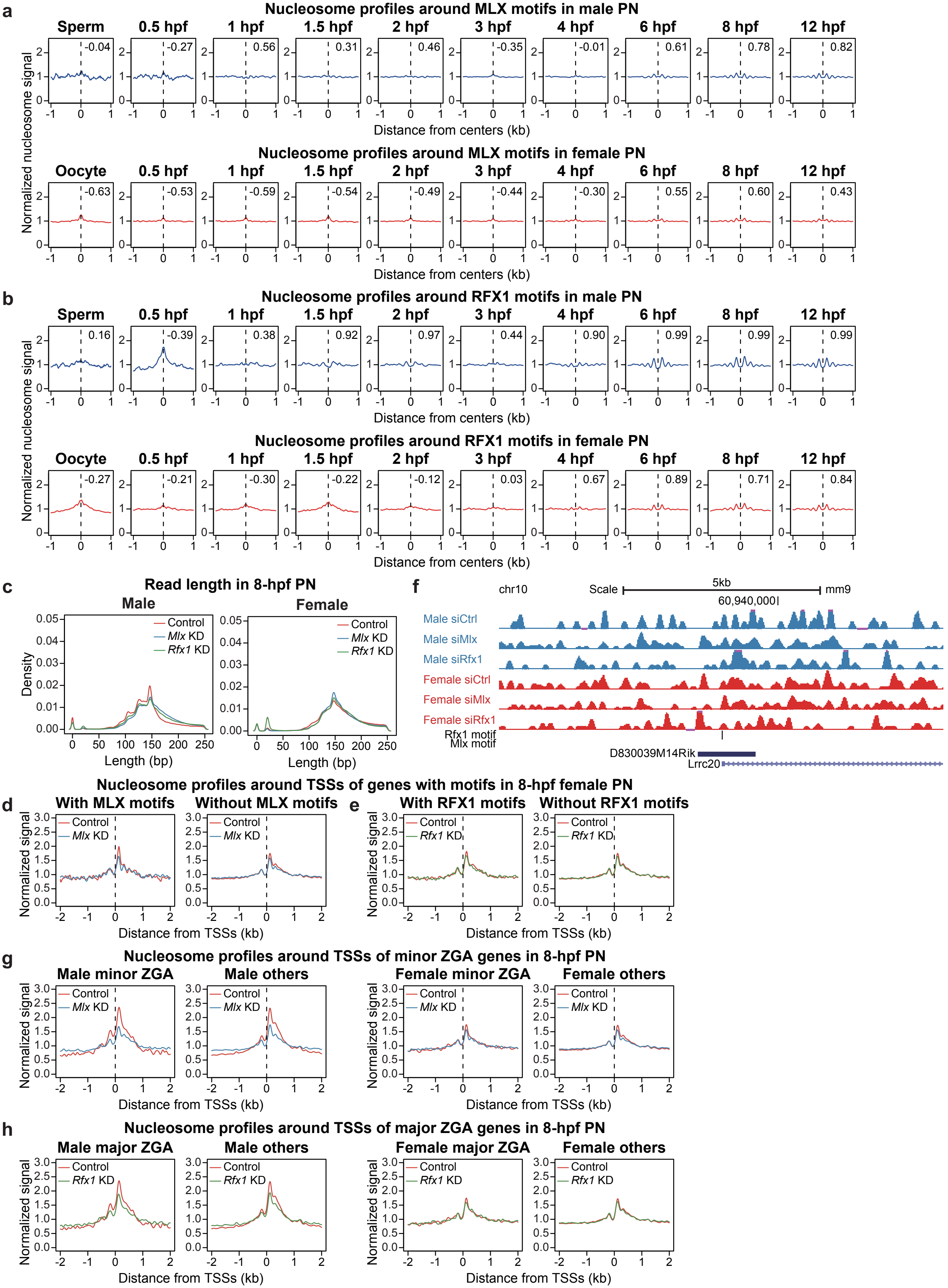
a
Nucleosome profiles around MLX motifs in male PN
Sperm
0.5 hpf
1 hpf
1.5 hpf
2 hpf
3 hpf
4 hpf
6 hpf
8 hpf
12 hpf
-0.04
-0.27
0.56
0.31
0.46
-0.35
-0.01
0.61
0.78
0.82
Normalized nucleosome signal
Distance from centers (kb)
Nucleosome profiles around MLX motifs in female PN
Oocyte
-0.63
-0.53
-0.59
-0.54
-0.49
-0.44
-0.30
0.55
0.60
0.43
b
Nucleosome profiles around RFX1 motifs in male PN
0.16
-0.39
0.38
0.92
0.97
0.44
0.90
0.99
0.99
0.99
Nucleosome profiles around RFX1 motifs in female PN
-0.27
-0.21
-0.30
-0.22
-0.12
0.03
0.67
0.89
0.71
0.84
c
Read length in 8-hpf PN
Male
Female
Control
Mlx KD
Rfx1 KD
Density
Length (bp)
f
chr10
Scale
5kb
mm9
60,940,000
Male siCtrl
Male siMlx
Male siRfx1
Female siCtrl
Female siMlx
Female siRfx1
Rfx1 motif
Mlx motif
D830039M14Rik
Lrrc20
Nucleosome profiles around TSSs of genes with motifs in 8-hpf female PN
d
With MLX motifs
Without MLX motifs
e
With RFX1 motifs
Without RFX1 motifs
Normalized signal
Distance from TSSs (kb)
g
Nucleosome profiles around TSSs of minor ZGA genes in 8-hpf PN
Male minor ZGA
Male others
Female minor ZGA
Female others
h
Nucleosome profiles around TSSs of major ZGA genes in 8-hpf PN
Male major ZGA
Male others
Female major ZGA
Female others

### Figure S10

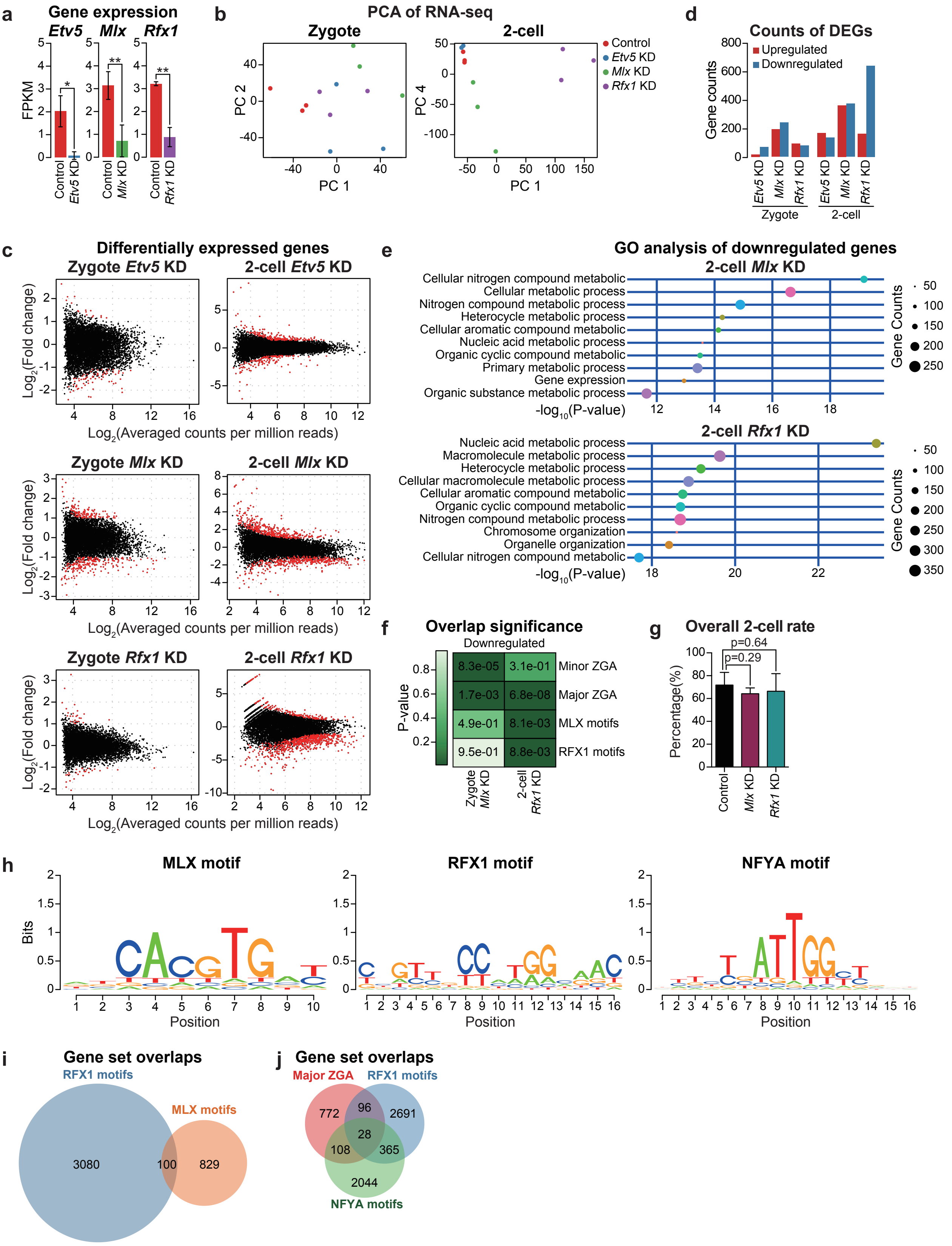
a
Gene expression
Etv5 Mlx Rfx1
FPKM
Control
Etv5 KD
Mlx KD
Rfx1 KD
*
**
**
b
PCA of RNA-seq
Zygote
2-cell
PC 1
PC 2
PC 4
Control
Etv5 KD
Mlx KD
Rfx1 KD
d
Counts of DEGs
Gene counts
Upregulated
Downregulated
Zygote
2-cell
c
Differentially expressed genes
Zygote Etv5 KD
2-cell Etv5 KD
Zygote Mlx KD
2-cell Mlx KD
Zygote Rfx1 KD
2-cell Rfx1 KD
Log2(Fold change)
Log2(Averaged counts per million reads)
e
GO analysis of downregulated genes
2-cell Mlx KD
Cellular nitrogen compound metabolic
Cellular metabolic process
Nitrogen compound metabolic process
Heterocycle metabolic process
Cellular aromatic compound metabolic
Nucleic acid metabolic process
Organic cyclic compound metabolic
Primary metabolic process
Gene expression
Organic substance metabolic process
-log10(P-value)
Gene Counts
2-cell Rfx1 KD
Nucleic acid metabolic process
Macromolecule metabolic process
Heterocycle metabolic process
Cellular macromolecule metabolic process
Cellular aromatic compound metabolic
Organic cyclic compound metabolic
Nitrogen compound metabolic process
Chromosome organization
Organelle organization
Cellular nitrogen compound metabolic
-log10(P-value)
Gene Counts
f
Overlap significance
Downregulated
P-value
8.3e-05 3.1e-01 Minor ZGA
1.7e-03 6.8e-08 Major ZGA
4.9e-01 8.1e-03 MLX motifs
9.5e-01 8.8e-03 RFX1 motifs
Zygote Mlx KD
2-cell Rfx1 KD
g
Overall 2-cell rate
p=0.64
p=0.29
Percentage(%)
Control
Mlx KD
Rfx1 KD
h
MLX motif
RFX1 motif
NFYA motif
Bits
Position
i
Gene set overlaps
RFX1 motifs
MLX motifs
3080
100
829
j
Gene set overlaps
Major ZGA
RFX1 motifs
NFYA motifs
772
96
2691
28
108
365
2044
